# Variable responses of alpine-plant communities to warming and loss of dominant species

**DOI:** 10.1101/2024.08.13.604445

**Authors:** ML Marraffini, NJ Sanders, MK Sundquist, AT Classen, JR Deslippe, JS He, JR McLaren, C Rixen, S Wipf, C Chisholm, J Giejsztowt, C Prager, DB Stouffer

**Affiliations:** Centre for Integrative Ecology, School of Biological Sciences, University of Canterbury, Christchurch, NZ; Department of Ecology and Evolutionary Biology, University of Michigan, Ann Arbor, MI 48109; Natural History Museum of Denmark, University of Copenhagen, Copenhagen, Denmark; The Gund Institute for Environment, University of Vermont, Burlington, VT, 05405; Department of Ecology and Environmental Sciences, Umeå University, SE901 87 Umeå Sweden; Centre for Biodiversity and Restoration Ecology, School of Biological Sciences, Victoria University of Wellington, Wellington, NZ; Institute of Ecology, College of Urban and Environmental Sciences, Peking University, Beijing, CN; Department of Biological Sciences, University of Texas at El Paso, El Paso, TX, USA; WSL Institute for Snow and Avalanche Research SLF, Flüelastrasse 11, CH-7260 Davos; Institute of Integrative Biology, Dept of Environmental Systems Science, ETH Zürich, CH-8092 Zürich, CH; Rubenstein School of Environment and Natural Resources, University of Vermont, Burlington, VT, USA

**Author notes:** **Author emails:** Sanders NJ,; Sundquist MK,; Classen AT,; Deslippe JR,; He JS,; McLaren JR,; Rixen C,; Wipf S,; Chisholm C; Giejsztowt J; Prager C; Stouffer DB. **Corresponding Author:** Michelle L. Marraffini. **Author Contributions** MLM and DBS conceived of the idea. NJS, MKS, ATC, JRD, JSH, JRM, and CR received funding for and coordinated field efforts as well as collected the data with participation from SW, CC, JG, and CP. MLM completed the data analysis and interpretation. MLM and DBS wrote the draft and all authors contributed to the final manuscript.

**Keywords:** Species interactions, Population dynamics, Global Change, Warming Experiment, Removal Experiment

## Abstract

Responses of ecological communities to perturbations are inherently variable because responses of their constituent populations also vary. Species within a single community may show combinations of no response, positive responses, and negative responses to any given perturbation often canceling each other out resulting in small or no signal that the community level. Here we explore the impacts of warming and loss of the dominant species on alpine ecosystems in a global study. We investigate warming and species-loss treatments on population- and community-level dynamics across alpine-plant communities at two elevations in five globally-distributed mountain locations. Communities showed varied responses to treatments; no community showed strong responses to a single treatment. Rather, most sites were influenced by both perturbations. Populations within these communities responded idiosyncratically, suggesting that constituent species are not all equally robust to perturbations even when community-level effects appear weak. Our results highlight the challenge of making general predictions about population- and community-level responses of alpine ecosystems in the face of present and future perturbations.

---

Environmental conditions and species interactions shape patterns of diversity and abundance under ambient conditions and influence community responds to perturbations. Species respond directly to changing environmental factors based on individual and population-level characteristics, such as physiological tolerances that limit their distribution (Grime 1979, Huston 1999, Pavoine *et al*. 2011). Additionally, species respond indirectly to environmental changes through shifts in the distribution, abundance, or behavior of their natural enemies and beneficial interaction partners (e.g. Dunson & Travis 1991, Davis *et al*. 1998, Sanford 1999, Hawkins *et al*. 2009). Although long recognized as biologically important, consequences of variation in environmental temperature have renewed importance as global climate change alters both mean and extreme temperatures (Jentsch *et al*. 2007, Smith 2011, Turner *et al*. 2020). While general patterns in response to global climate change may be predictable, making accurate predictions for specific species or ecosystems is complicated by the interplay of direct and indirect effects (Kordas *et al*. 2011).

Species interactions are also critical drivers of patterns in diversity and abundance across landscapes (Grime 1979, May & McLean 2007, Bruno *et al*. 2003, Klein *et al*. 2004, Molau 2010). Dominant species can exert significant influence by affecting the interactions among other species, a concept partially explained by the ‘mass ratio’ hypothesis, which posits that ecosystem functioning is largely determined by the traits of the most abundant species (Weaver & Clements 1938, Dayton 1972, Paine 1974, 1969, Grime 1998, Ellison *et al*. 2005). For example, *Empetrum nigrum* subsp. *hermaphroditum*, an evergreen dwarf shrub which dominates low-nutrient alpine ecosystems (Tybirk *et al*. 2000), impacts other species by forming dense mats and producing allelopathic compounds (Nilsson 1994). This in turn reduces germination and survival of co-occurring species, leading to reduced vascular plant richness (Aerts 2010). Given the breadth of potential impacts, declines in the abundance or complete loss of a dominant species can have cascading effects throughout a community.

While there is an ever-growing body of literature that demonstrates the effects of warming temperatures (Wolkovich *et al*. 2012, Sundqvist *et al*. 2013) or species interactions (Adler *et al*. 2007, Brooker *et al*. 2008) on population- and community-dynamics, examining both drivers concurrently is essential to understanding how communities coexist and respond to future environmental changes (Matías *et al*. 2018, Bimler *et al*. 2018). Change in an environmental variable could alter abundance of a species indirectly, as interactions can vary in outcome and strength within populations and across environmental gradients (Bowen 1980, Thompson 1988, Adler *et al*. 2012, Bimler *et al*. 2018). For example, the effects of *E. nigrum* are found to be dependent on environmental variables including soil moisture and geomorphic disturbance (Mod *et al*. 2014). Similarly, work in annual-plant communities has shown that interactions between plants may shift from competitive to facilitative along environmental gradients (He *et al*. 2013, Badano *et al*. 2007, Bimler *et al*. 2018). This aligns with the Stress Gradient Hypothesis, which suggests that facilitative interactions become more common in more stressful environments, while competitive interactions dominate in less stressful conditions Bertness & Callaway (1994), Callaway (1995), He *et al*. (2013). Key interactions, that are sensitive to temperature, can act as “leverage points” amplifying small changes in climate into large consequences for the community (Sanford 1999). This indirect response to the environment stems from species-specific responses, which can change the absolute or relative abundance of interaction partners or relative strength of interactions themselves (Adler *et al*. 2009, Bimler *et al*. 2018). As a result, the interplay of environmental conditions and species interactions remains an important piece of the puzzle, as modified species interactions can lead to differing conclusions about dynamics and stability of whole communities (Suttle *et al*. 2007).

Previous studies have explored variation in plant responses to experimental warming and composition manipulation with mixed results (e.g. Hobbie *et al*. 1999, Rixen & Mulder 2009, Cavieres & Sierra-Almeida 2012) because the scales (community vs population) measured among experiments varied. One way forward is to measure population- and community-level responses to these drivers, which represent direct and indirect effects of climate change, through coordinated experiments at multiple sites. Therefore, we examine the effects of warming and species removal treatments on plant cover, in alpine communities across two elevations (high and low) in a global experimental study. Elevation gradients can serve as natural experiments for how communities and ecosystems respond to changes in climate as they represent a combination of co-varying abiotic factors, including temperature and soil moisture (Sundqvist *et al*. 2013, McCain & Colwell 2011, Pauli *et al*. 2014). High elevation plant communities are predicted to be strongly regulated by environmental factors (Grime 1977) since they are often subject to more physically stressful conditions than low elevation plant communities (Callaway 1995). We use experimental treatments (warming and species removal) as alternative hypotheses of the relative importance of abiotic and biotic drivers in determining percent cover of alpine-plants in five locations over four to five years. We expected that the effect of species removal would be greater than the direct effect of warming, as these communities may experience a range of temperatures and are long-lived, allowing them to buffer against extremes. High elevations are temperature limited while intensity of competitive interactions is greater at low elevations. Therefore, we expect the impact of species removal to be greater at low elevations. Additionally, we expect variation among locations in the effects of warming and removal due to inherent differences and co-factors such as water availability.

## Methods

### Field Methods

We conducted replicated field experiments at high and low elevation sites at five locations: Canada (CA), China (CN), New Zealand (NZ), Switzerland (CH), and United States (US) (Table S1). Henceforth, we will refer to specific elevations within a location as “sites”. At each site, the experiment consisted of eight replicate blocks, containing 2 m × 2 m plots with 2 m buffers. Plots were randomly assigned to one of four treatments: ambient control (ambient, A), ambient temperature with dominant removal (removal, R), warming without removal (warming, W), and warming and dominant removal (warming and removal, R:W). This yielded 32 total plots per site. Removal treatments consisted of above-ground biomass removal of a single locally dominant species, repeated annually, by clipping to ground level (and, as necessary, herbicide application). The dominant species was the same for all plots at a given site but varied across locations (CA, US, CN, NZ, CH) (Table S2 and S1.1), and were not included in the analyses described below. Warming plots consisted of open top chambers covered with hexagonal polycarbonate, sloping transparent sides, and an internal diameter of 1.5 m. This stayed in place during the growing season and achieved an increase of 1-3 °C (Henry & Molau 1997, Molau & Mølgaard 1996) (Fig S1). For the warming and removal treatment, removal took place over the 2 m × 2 m plot while the warming encompassed 1.5 m diameter within that. During the growing season, annual sampling within each plot consisted of visual percent-cover estimates to monitor changes in individual species cover. US and CH were sampled over five years, while the remaining three sites were samples over four years. However, the CN treatment plots were not measured for all years (i.e. some were measured in years 1, 2, and 3 but not year 4, or other combinations), leading to lower replication at this site. All plants were recorded to the lowest taxonomic unit with laboratory identifications as needed (Table S5).

### Population-Dynamics Model

We developed a discrete-time population-dynamics model to predict year-to-year changes in percent cover of these plants as a combination of density-independent and density-dependent changes (Rees *et al*. 1996, Adler *et al*. 2012, Martorell & Freckleton 2014). We designed our models such that they would enable us to separate community-wide (experimental plot-level) responses from species-specific responses by estimating a grand mean parameter which represents the ‘average’ species in that community and individual species deviations from this grand mean. Specifically, we estimated percent cover *N_i,t_* of a focal species *i* in year *t* as a function of the prior year’s percent cover *N_i,t−_*_1_ with a model that takes the general form:

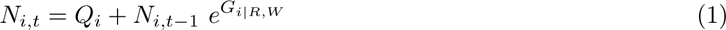

*Q_i_* represents density-independent increase in percent cover (influx), which may be attributed to recruitment, lateral encroachment into plots, and/or plants emerging from seed. Note that this *Q_i_* term is not influenced by previous percent cover and allows us to account for observations of a species that was unobserved in the previous year, which occurred in 42% of our observations (see Supplemental). Since experimental plots were randomly assigned throughout the landscape, we assumed that this density-independent influx was unaffected by treatments. In contrast, the model component *N_i,t−_*_1_*e^G_i|R,W_^* captured density-dependent changes in percent cover since it is multiplied by previous percent cover; these changes can be driven, for example, by growth or mortality.

Density-independent change in percent cover, *Q_i_* of a species *i* is given by the form:

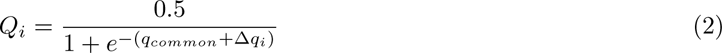

Hence, *Q_i_* is constrained with 0.5 as a maximum to be more biologically plausible and improve model fit. The combination of the community-wide grand mean (*q_common_*) and the species-specific deviations (Δ*q_i_*) can be interpreted as the inferred density-independent change in percent cover for that species.

We likewise parameterized density-dependent change *G_i|R,W_* of a species *i* such that it contained both the common (*g*_common_*_|R,W_*) and species-specific (Δ*g_i|R,W_*) effects, but here we allowed these effects to also vary as a consequence of treatment. Mathematically, this takes the form:

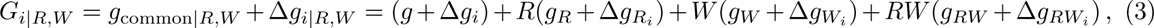

where the parameters *R* (removal) and *W* (warming) equal 1 in plots where that treatment was applied and otherwise equal 0. Species-specific responses for each species *i* under each treatment were captured as devia-tions from grand means via the parameters Δ*g_i_*, Δ*g_Ri_*, Δ*g_Wi_*, and Δ*g_RWi_*. The combination of community-wide grand mean and species-specific deviations can be interpreted as the inferred density-dependent change in percent cover for that species. For example, the estimate of *g*_all_*_|R_* = *g* + *g_R_* can be interpreted as the component of density-dependent change of the ‘average’ plant at a given site under the removal treatment while the estimate of Δ*g_i|R_* = Δ*g_i_* + Δ*g_Ri_* indicates how species *i* deviates from this grand mean.

### Model Inference

Since our main goal is to identify the importance of treatments to change in percent cover of individual alpine-plants, we chose to investigate each site (location × elevation) separately. Community composition varied both between elevations at the same location and across locations, further supporting this decision. We concentrated our analysis on widespread species within each site (Figs S3 and S4), determined as species observed in at least 25% of total plots. For example, 0.25 × 32 plots × 4 years = 32 plots, which becomes the minimum number of plots a species must be present in to be analyzed (Table S1). While these widespread species represent a subset of the total species observed, they account for the majority of percent cover observed (see Supplemental S1.2, Table S1, Figs S14 and S15).

We used a Bayesian hierarchical model with Hamilton Markov Chain Monte Carlo (HMCMC) methods to infer the parameter values of our model (Eqs 1–3). We used a zero-inflated beta distribution and weakly informative priors to simulate predicted percent cover (data scaled to 0-1). We fit delta parameters as random effects for individual species, in order to estimate how each species responded to the treatments. This helped provide estimates for each species that are pulled towards the grand mean if its sample size is low (McElreath 2016). We performed sampling to determine the posterior distributions of model parameters with the function “brm” from the package “brms” (Bürkner 2017) in R (version 3.4.2) (R Core Team 2013). We ran two chains with a warm-up of 1000 iterations and 4000 sampling iterations each, for a final combined posterior of 6000 MCMC samples for each model. We determined that parameters converged when trace plots were well mixed and stationary, and when the Gelman-Rubin convergence diagnostic was close to one (Gelman & Rubin 1992, Brooks & Gelman 1998). See Supplemental for more specifics.

### Model Comparison

At each site (location × elevation), we compared six models to determine the relative importance of treatments in explaining variation in population dynamics (Table 1). Here, our models serve as alternative hypotheses; therefore, our main motivation is to determine which model has the best predictive accuracy, and is expected to fit future observations well, rather than assessing which model best matches the observed data (Aho *et al*. 2014). The Null model (*N_i,t_* = *N_i,t−_*_1_) predicts that percent cover is equal to previous percent cover (*N_i,t−_*_1_, if *Q_i_* and *G_i_* are zero) and acts as a well-defined baseline for comparison. The Recruitment model modifies this to include density-independent change in percent cover (influx, *Q_i_*), resulting in *N_i,t_* = *Q_i_* + *N_i,t−_*_1_. The Ambient model measures density-independent change in percent cover (*Q_i_*) and density-dependent change in percent cover in absence of any treatment effects (*N_i,t−_*_1_*e*^*g*+Δ*g_i_*^). Building on this, the single-treatment models (Removal or Warming) modify the density-dependent component by adding the main effect of the corresponding treatment (Table 1). The Removal + Warming model includes main effects for both treatments (Table 1); and finally, the Removal × Warming model contains the main effect of both treatments and their statistical interaction (Table 1).

**Table 1:** Model names and their density-dependent components which we compared for each site. Model parameters are as outlined in Eq (3)

| Model name | Density-dependent change in percent cover, $G_{i\vert R,W}$ |
| --- | --- |
| Null | 0 |
| Recruitment | 0 |
| Ambient | $g + \Delta g_i$ |
| Removal | $g + \Delta g_i + R(g_R + \Delta g_{R_i})$ |
| Warming | $g + \Delta g_i + W(g_W + \Delta g_{W_i})$ |
| Removal + Warming | $g + \Delta g_i + R(g_R + \Delta g_{R_i}) + W(g_W + \Delta g_{W_i})$ |
| Removal × Warming | $g + \Delta g_i + R(g_R + \Delta g_{R_i}) + W(g_W + \Delta g_{W_i}) + RW(g_{RW} + \Delta g_{RW_i})$ |

We used the Widely Applicable Information Criteria (WAIC) to determine the best model or models for each site (Watanabe 2010, Burnham & Anderson 1998). We also used Akaike weight (based on WAIC) to compare the relative predictive accuracy of models, where larger values indicate greater support (Burnham & Anderson 2002, McElreath 2016). We defined best-fit models as those with the lowest WAIC and an Akaike weight greater than 0.8; when a single model did not meet this criteria, we used model averaging to combine models’ posterior distributions based on Akaike weights.

## Results

### Community-Level Results

While we fit our model to observations of individual plant species, we can scale up to get a sense of what is happening at plot-level. Here 70% (664 out of 946) of plot-level predicted percent cover were within credible intervals (Fig 1). Additionally, examining the grand mean of a community gives a picture of how the ‘average’ species responded to treatments, but as we will see later, these communities are made up of many species that themselves respond to treatments idiosyncratically.

**Figure 1:**
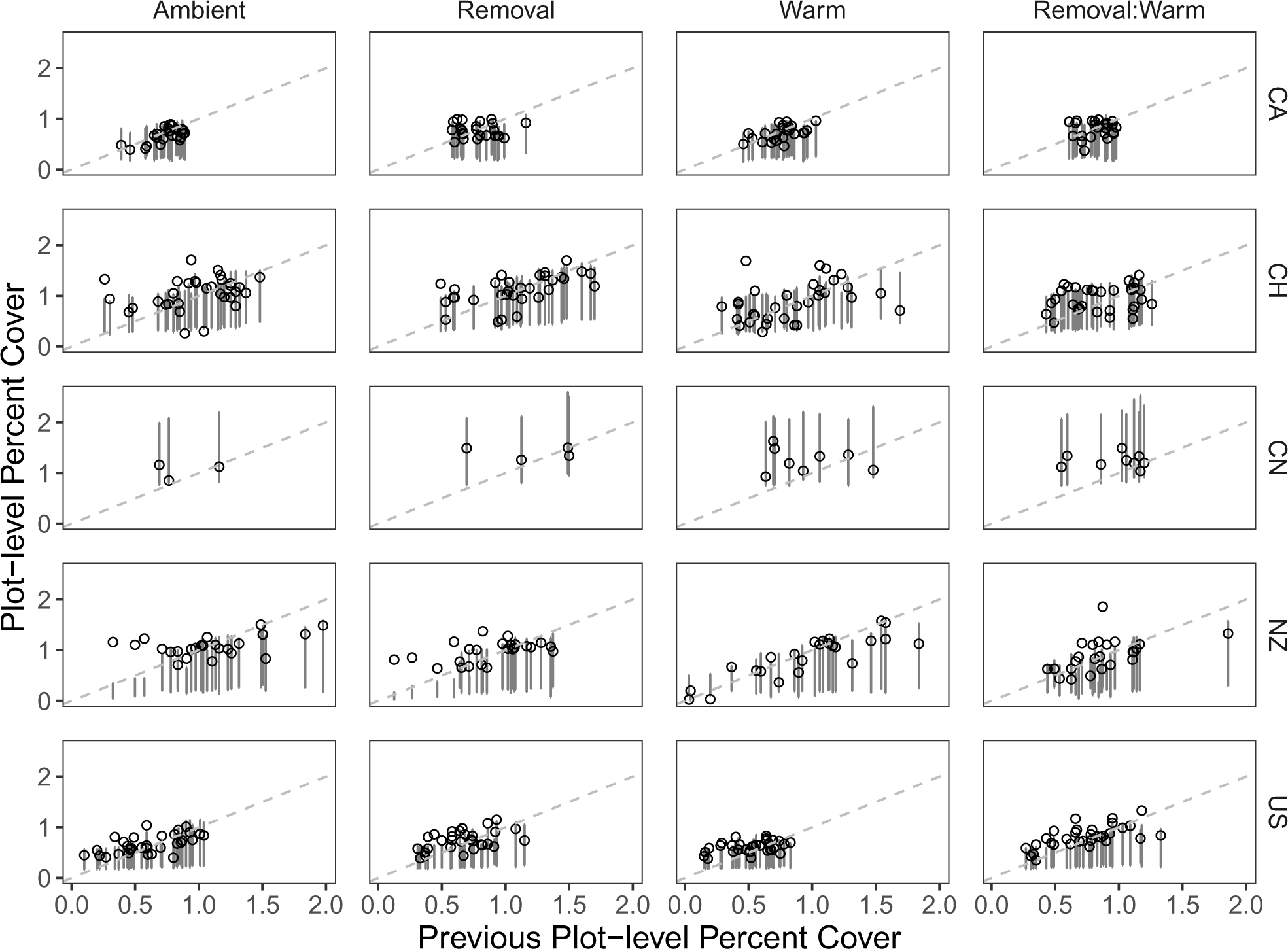
Year-to-year total percent cover for all plots at low elevation across all sites. Points show the total observed percent cover in a plot while vertical lines show the credible interval (89%) of our statistical model’s predicted total percent cover for that plot. Species-specific predicted percent cover is calculated according to Eqn 1 then summed across all species within a plot to yield predicted plot-level percent cover. As a visual guide, the dashed line represents the 1:1 or where previous percent cover would equal current percent cover (i.e. when total percent cover in a plot neither increases nor decreases). Comparable plots for high elevation sites can be found in the Supplemental (Fig S5). Data scaled for beta distribution (i.e. between 0-1 for each species).

Best-fit models varied among locations (CA, CH, CN, NZ, and US) and elevation (High vs. Low) within a location, highlighting the complicated nature of communities, especially in responses to multiple perturbations. Three sites (high elevation CH, low elevation NZ, and low elevation US) showed no obvious effect of treatments with the Ambient model receiving all the WAIC support and Akaike weight (Table 2). Four sites (low elevation CA, low elevation CH, high elevation NZ, and high elevation US) showed support for treatment effects with a treatment model gaining *>* 50% of WAIC weight (Table 2). While no site showed exclusive support for the Warming model, multiple sites showed marginal support for including this treatment based on Akaike weights (high elevation CA, high elevation CN, high elevation NZ, and high elevation US, Table 2). Specifically, the Warming model accounted for 70% of the Akaike weights in high elevation US model comparison (missing our 80% cutoff for a winning model, Table 2) and did have a small positive effect on the community (Table 3). Similarly, Removal model received 64% of Akaike weight at low elevation CA (Table 2). Additionally, multiple sites showed some support for the interaction treatment (Removal × Warming), with this model receiving approximately 5% WAIC weight (except high elevation CN, see Supplemental). Both low elevation CH and high elevation NZ showed some support for both treatments. High elevation NZ showed additional support for the warming × removal treatment model. While variable in nature, this suggests that our models are capable of capturing a community-level percent cover response to treatments when it is present.

**Table 2:**
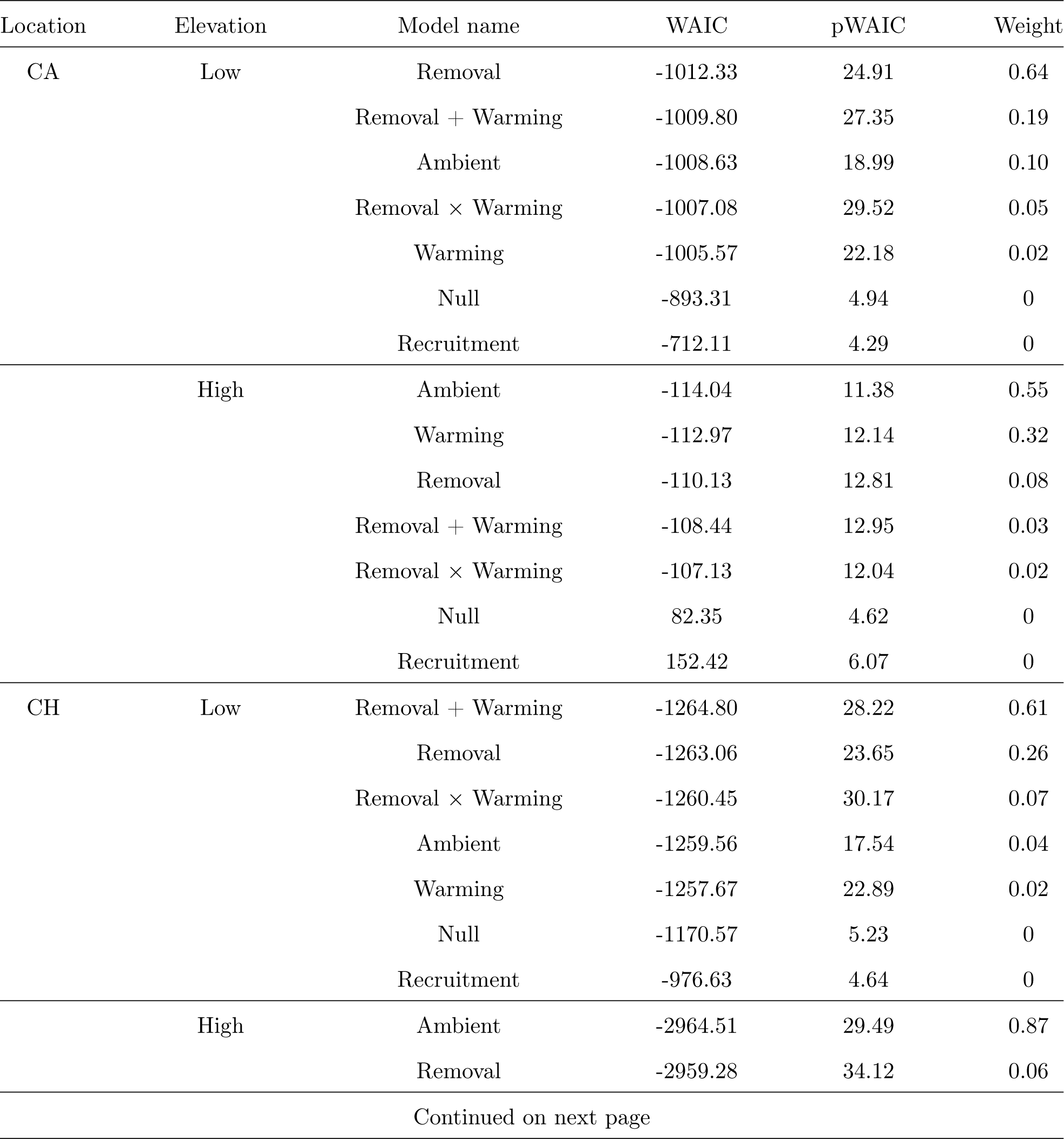
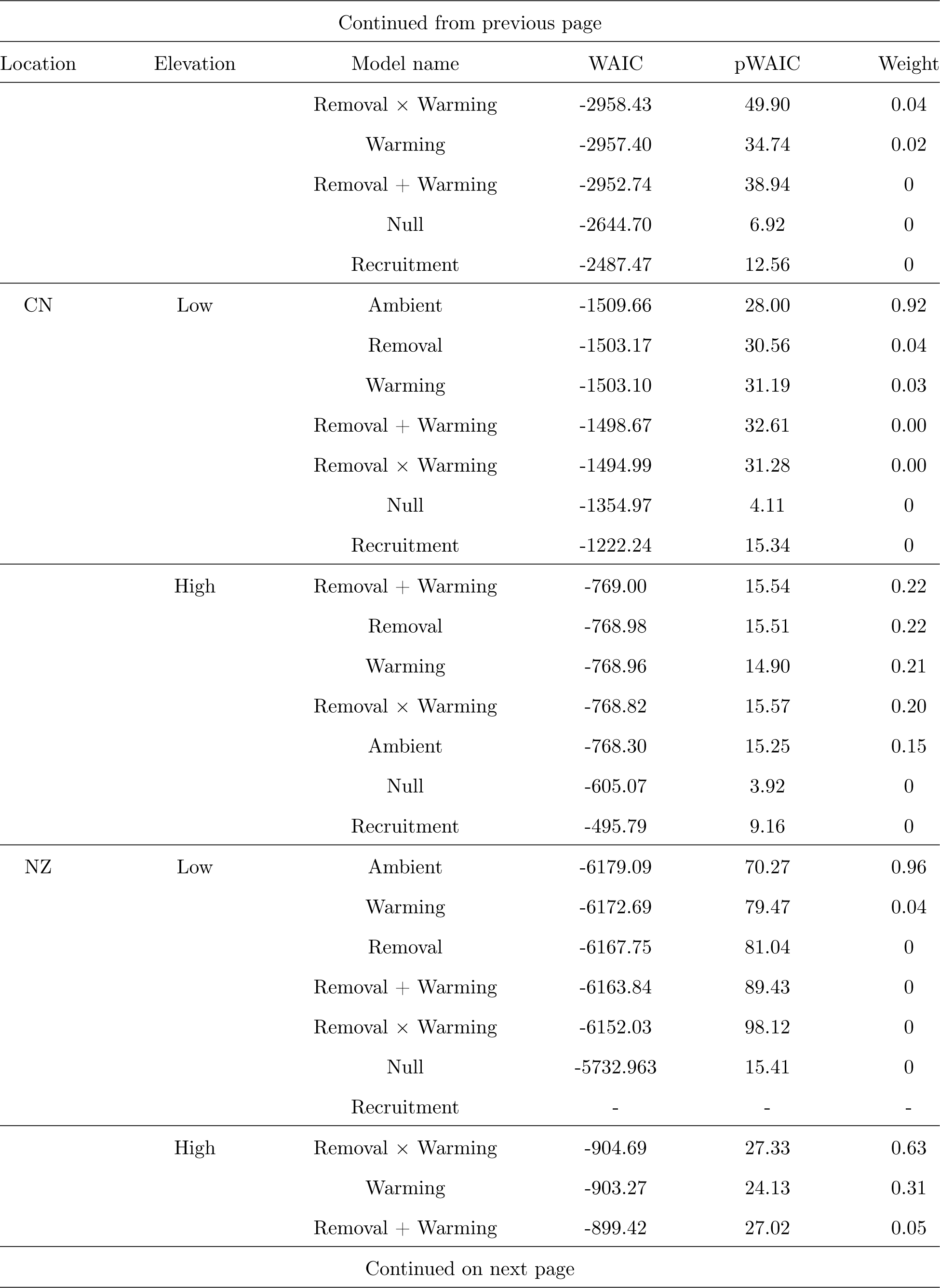
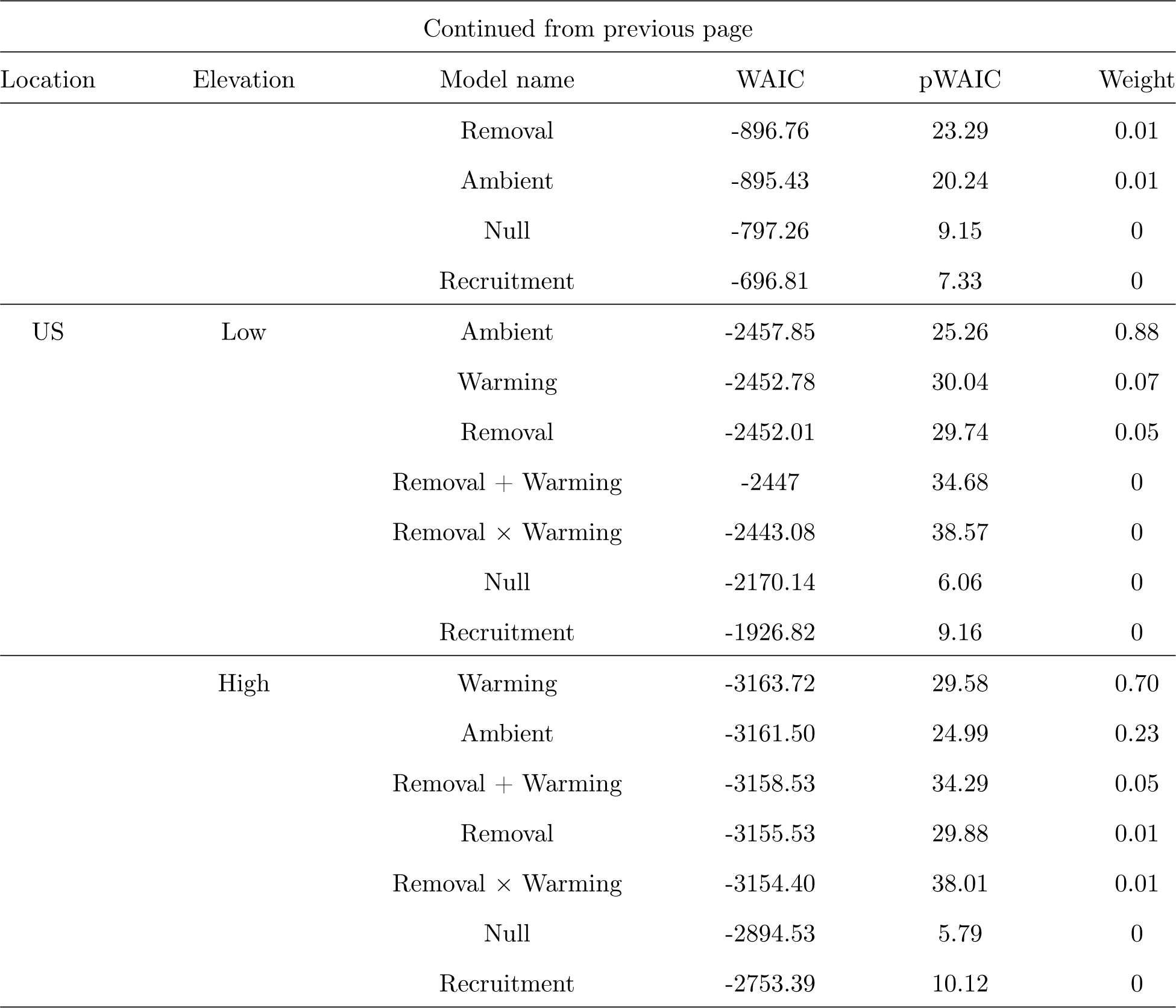
Model comparison table giving information criteria for each model tested for low and high elevation communities in Canada (CA), Switzerland (CH), China (CN), New Zealand (NZ), and the United States (US). WAIC (Widely Applicable Information Criteria) penalizes models for parameters and the lowest WAIC reflects the best-fit model. pWAIC is the effective number of parameters and provides information on how flexible each model is in fitting the sample. Weight refers to Akaike Weight for each model, interpreted as an estimate of the probability that the model will make the best predictions of new data based on the the set of models considered. The low elevation NZ recruitment only model did not converge.

**Table 3:** Median estimates from the posterior distribution of model parameters. Note that these are un-transformed and hence correspond to Eq 2 and 3. Composite models based on Akaike weights do not have Rhat values.

| Location | Elevation | Model Name | Parameter | Estimate | Lower 0.95 | Upper 0.95 | Rhat |
| --- | --- | --- | --- | --- | --- | --- | --- |
| CA | Low | Composite | $q$ | -2.33 | -2.44 | -2.21 | - |
| | | | $g$ | -1.20 | -1.36 | -1.03 | - |
| | | | $g_R$ | 0.05 | -0.13 | 0.22 | - |
| | | | $g_W$ | -0.07 | -0.18 | 0.03 | - |
| CA | High | Composite | $q$ | -2.18 | -2.58 | -1.83 | - |
| | | | $g$ | -0.34 | -0.70 | 0.02 | - |
| | | | $g_R$ | -0.04 | -0.31 | 0.24 | - |
| | | | $g_W$ | -0.24 | -0.51 | 0.03 | - |
| CH | Low | Composite | $q$ | -1.812 | -1.94 | -1.68 | - |
| | | | $g$ | -0.71 | -1.02 | -0.44 | - |
| | | | $g_R$ | 0.04 | -0.05 | 0.24 | - |
| | | | $g_W$ | -0.04 | -0.14 | 0.05 | - |
| | | | $g_{RW}$ | 0.15 | 0.01 | 0.31 | - |
| CH | High | Ambient | $q$ | -2.66 | -2.93 | -2.35 | 1.01 |
| | | | $g$ | -1.02 | -1.42 | -0.64 | 1 |
| CN | Low | Composite | $q$ | -1.81 | -1.94 | -1.67 | - |
| | | | $g$ | -0.79 | -1.02 | -0.60 | - |
| | | | $g_R$ | 0.14 | 0.02 | 0.26 | - |
| CN | High | Composite | $q$ | -2.66 | -2.76 | -2.57 | - |
| | | | $g$ | -1.01 | 1.14 | -0.88 | - |
| | | | $g_R$ | -0.09 | -0.14 | -0.04 | - |
| | | | $g_W$ | -0.005 | -0.05 | 0.05 | - |
| | | | $g_{RW}$ | 0.06 | -0.05 | 0.18 | - |
| NZ | Low | Ambient | $q$ | -3.93 | -4.29 | -3.58 | 1 |
| | | | $g$ | -0.63 | -0.87 | -0.41 | 1 |
| NZ | High | Composite | $q$ | -3.70 | -3.93 | -3.49 | - |
| | | | $g$ | -0.62 | -0.84 | -0.42 | - |
| | | | $g_R$ | 0.04 | -0.09 | 0.18 | - |
| | | | $g_W$ | -0.04 | -0.16 | 0.11 | - |
| | | | $g_{RW}$ | -0.26 | -0.47 | -0.02 | - |
| US | Low | Ambient | $q$ | -2.61 | -2.90 | -2.33 | 1 |
| | | | $g$ | -1.12 | -1.52 | -0.63 | 1 |
| US | High | Composite | $q$ | -2.32 | -2.43 | -2.21 | - |
| | | | $g$ | -1.05 | -1.16 | -0.95 | - |
| | | | $g_W$ | 0.18 | 0.12 | 0.26 | - |

To envision how our models predict percent cover, we take an example of low elevation CA. If the ‘average’ species had a previous percent cover of 10% (0.10 on our scale from 0 to 1), the model estimates its percent cover in the following year to be 7.44% in the ambient plots, 7.59% in removal plots, and 7.24% in Warming plots. This median estimate is made up of both the density-independent and density-dependent factors. Density-independent influx alone predicts the next year’s percent cover to be 4.43%. While, the median estimate of density-dependent change in percent cover in the ambient treatment is *g* = *−*1.20 (Table 3). To understand the meaning of this value, we take the exponential and then multiply it by previous percent cover (*N_t−_*_1_ *× e^g_common_^* = 10% *× e^−^*^1.20^ = 3.01%). At this site, multiple treatment models received support (Table 3). In the removal treatment, the median density-dependent change (*g_common_* + *g_Ri_*) in percent cover was estimated as *N_t−_*_1_ *×e^g_common_^* = 10% *×e*^(^*^−^*^1.20+0.05)^ = 3.16% as this treatment adds to the value from the ambient estimate. This suggests that removal of the dominant species slightly ‘helped’ the average species in this community compared to the ambient control. In contrast, the Warming treatment lowered percent cover of the average species compared to ambient plots suggesting that species in this community tend to be vulnerable to the direct effects of Warming (*N_t−_*_1_ *× e^g_common_^* = 10% *× e*^(^*^−^*^1.20^*^−^*^0.07)^ = 2.81%). Consider high elevation CH as another example. If the average species there had a previous percent cover of 10% (0.10), the model estimates its median percent cover in the following year to be 6.87% in the ambient plots. This predicted percent cover is made up of density-independent influx (*Q_common_*) of 3.26% and a density-dependent component of 3.61% with this site showing support for the ambient only model. If the average species instead had a previous percent cover of 1%, the model estimates its abundance would increase in the following year to 3.63% in all the plots.

While there was support for a measurable influx of new species into plots, our model distinguished this from how species already present were responding to treatments. At the community-level, median estimates of year-to-year density-independent influx of species varied by site ranging from *q_common_* = *−*3.93 to *q_common_* = *−*1.81 amounting to 0.95% and 7.03% cover respectively. Species in low elevation NZ received the lowest average contribution of this influx to variation in species percent cover and low elevation CH received the highest contributions (Table 3). Density-independent influx may have resulted from an influx of seeds, germination of seeds from the seed bank, or lateral encroachment of neighboring plants which may be allowing plots to maintain percent cover in-spite of density-dependent change in percent cover (*e^g_common_^*) of less than 1 in some areas.

### Species-Level Results

Communities in this study were composed of plant species that varied in initial percent cover and also in their density-dependent change in percent cover and responses to treatments. Though the effects of treatments were not always apparent at a community-level scale (Fig 2; Figs S6-S14), our models were able to detect impacts at the species-specific scale. Indeed, some of the statistical support for including or removing different treatment effects may be due to differences across species in addition to any signal that was detected at the community scale. This notion is supported further by the observation that species on the whole responded idiosyncratically to treatments within each site. That is, there were species in each site that exhibited responses similar to, more exaggerated than, or contrary to the grand mean density-dependent change in percent cover (Fig 2; Figs S6-S14). There were no overarching patterns in the way species responded and we did not detect any strong responses of functional groups (see Supplemental, Table S5).

**Figure 2:**
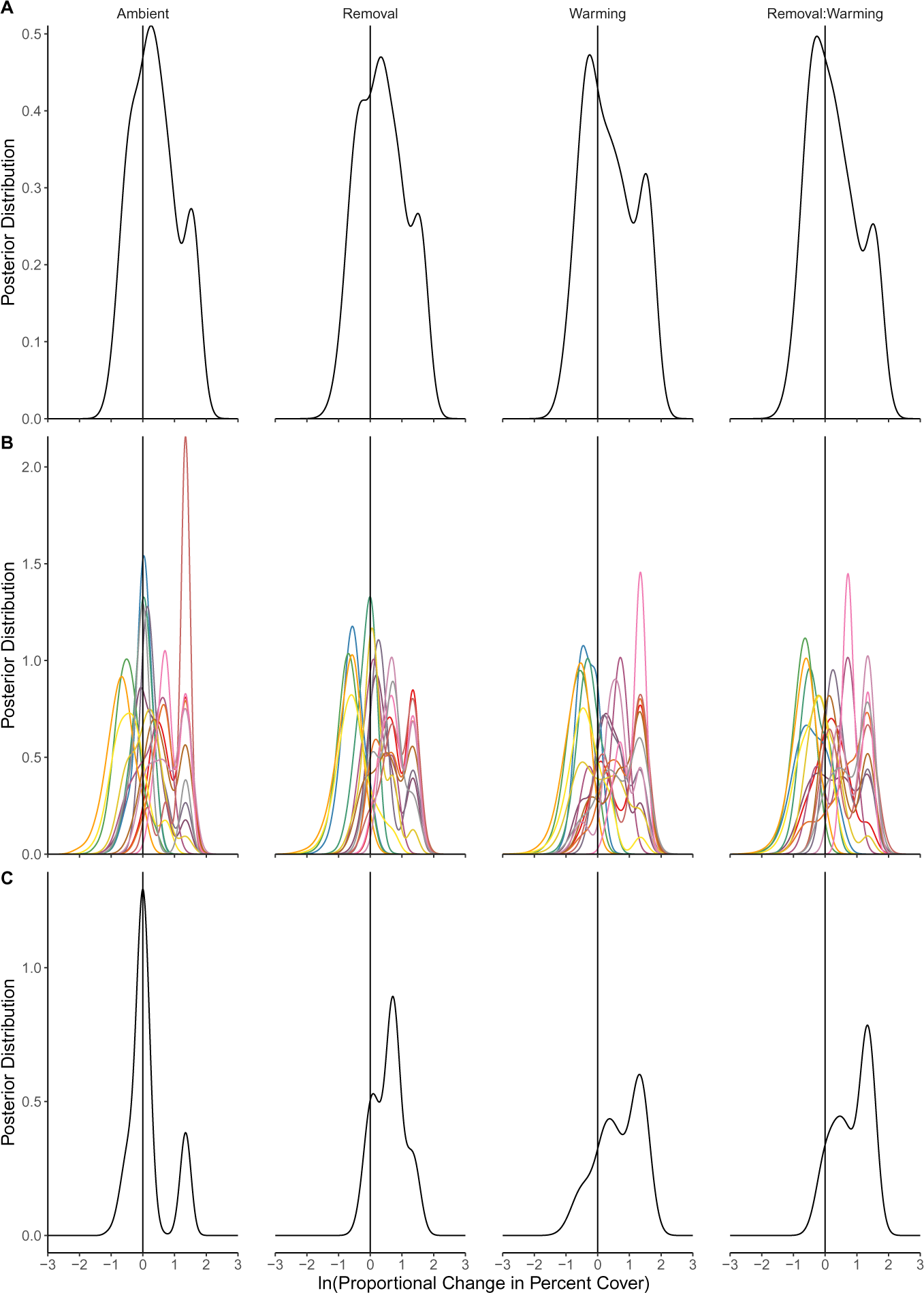
Posterior distribution of predicted logarithmic proportional change in cover within the low elevation Canada site based on the predictions of the best-fit model. This site acts as a representative example of the variable responses to treatment we observe at the species-specific level. Logarithmic proportional change in cover is calculated as (ln(*N_i,t_/N_i,t−_*_1_) = ln(*Q_i_/N_i,t−_*_1_ + *e^Gi|RW^*)) as sampled from the joint posterior for parameters in our statistical model. A logarithmic change of zero (ln(1) = 0; solid vertical line) represents no change over time suggesting that populations are at/near their equilibrium. A) Shows the posterior predictions based on the community-level grand mean or the ‘average’ species, which in this case only shows minor effects of the removal treatment. B) Shows the posterior predictions for all species at this site illustrating how species vary within and among treatments. C) Shows the changes in a single species *Veronica wormskjoldii*. This species showed an increase in density-dependent change in cover with the removal of the dominant species compared to the values near zero under ambient conditions. Warming and the Removal × Warming treatments did not receive any model support at this site so their predictions reflect the ambient and removal predictions, respectively. Since this metric includes observed previous percent cover, differences seen in unsupported treatments reflect variations in species percent cover rather than inferred parameters.

For example, at low elevation CA, community-level grand means showed support for the inclusion of removal treatment but with a small effect on grand mean density-dependent change in percent cover (Fig 2). Within this site, species showed variable logarithmic proportional change in cover (ln(*N_i,t_/N_i,t−_*_1_)) under ambient conditions and the same species showed different yet still variable changes in percent cover after removal of the dominant species (Fig 2). Under ambient conditions, six species showed a mean change in percent cover of near zero suggesting that they remained close to their previous percent cover and are near equilibrium. After removal of the dominant species, an additional species showed no proportional change in their percent cover and remained at equilibrium. Three species that were near equilibrium under ambient conditions experienced a positive change in percent cover after the removal of the dominant. This suggests that some species increased their density-dependent change in percent cover in absence of the dominant compared to their growth under ambient conditions (Fig 2). On the other hand, one species showed a decrease in percent cover moving away from equilibrium. *Veronica wormskjoldii* showed an increase in median percent change in cover with the removal of the dominant species compared to ambient conditions (Fig 2C), differing from the average species proportional change in cover seen at this site (Fig 2A). Warming treatment at low elevation CA had similarly variable results with only one species showing no change. Overall species increased (4), decreased (3), or showed no change (9) compared to ambient conditions. This site acts as just one example of the different dynamics observed at the species-specific level (see Figs S6-S14).

## Discussion

Here, we explored the relative effects of warming and species-removal on alpine-plant communities using a combination of empirical data and population-dynamics modeling. We found that most communities did not show consistent responses to our experimental manipulations (Table 3), and percent cover of these communities remained relatively constant (Figs 1 and Fig S5) suggesting overall “resistance” after 4-5 years of treatments at the community level. This pattern of no/weak treatment effects was seen in both our population-dynamics model (Fig 1) and as well as a supplemental linear analysis (Table S3 and Fig S4). The relative importance of dominant species removal and warming temperatures varied among communities, with four out of ten communities showing moderate support (low elevation CA, low elevation CH, high elevation NZ, high elevation US) for treatments with varying inferred effects of warming and removal. While no community showed strong support for a single treatment as a predictor of change in percent cover, high elevation US showed limited support for the effects of warming, and low elevation CA showed limited support for removal of the dominant (Table 3). Beyond these general patterns, species within a community showed variable responses to perturbations, as some responded similarly to the ‘average’ species at their site while others deviated strongly from this average (Fig 2). These species-specific responses, possibly due to varying life-history strategies (i.e. forb compared to C3 graminoid) or soil specific effects (Ford & HilleRisLambers 2020), may allow a community to persist overall in the face of environmental change even as some individual populations decline (Pauli *et al*. 2014). However, we did not detect a pattern in the response of plant growth forms (see Supplemental, Table S5). The dominant species at each site come from a variety of functional groups (e.g. C3 graminoids, herbs, and woody shrubs) which did not dictate the remaining species’ response to the loss. Additionally, functional diversity of the remaining species also did not appear to dictate the importance of treatments or response to treatments (see Supplemental, Tables S4 and S5). Our results highlight the complicated nature of uncovering how drivers of population dynamics will be altered by the composition of the community and scale of the observation (community-vs. population-level; single-vs. multi-site) as well as by future and on-going disturbances such as climate change.

While we did not detect a shift in the importance of neighbors (i.e. the dominant species) through our model comparisons, we can see a shift in the influence of the dominant species among elevations which is consistent with the stress-gradient hypothesis (Bertness & Callaway 1994, Callaway 1995, He *et al*. 2013). For example, removal was somewhat important at the low elevation in Canada (64% of WAIC weight) but less so at the high elevation there (8% of WAIC weight). Here, we inferred higher percent cover (less density-dependent decrease) in removal compared to ambient treatments in low elevation communities suggesting previously negative (competitive) interactions with the dominant. High elevation CA communities on the other hand showed a decrease in percent cover with the loss of the dominant suggesting the dominant has facilitative effects. Similarly, low elevation China communities increased in percent cover while high elevation communities decreased in response to removal, though again with limited support for the removal treatment model. Small changes in total percent cover at the community level did not imply that these communities were robust to perturbations. Indeed, responses of the subdominant species were mixed suggesting that the dominant species, while influential, did not interact with all species equally or in the same manner (Fig 2). For example, low elevation Canada communities contained some species that were sensitive to and others that were resistant to the removal of the dominant species. On the other hand, high elevation Canada communities contained species that were more resistant to the removal of the dominant species. Previous work highlights that even small changes in the relative biomass of abundant species can cascade into large effects on ecosystem functioning (Grime 1998, Gaston & Fuller 2008). Thus even with limited community-level effects (small changes in percent cover) at some sites, the loss of the dominant species could lead to a shift in ecosystem functioning because remaining species vary in percent cover resulting in changes in above- and below-ground primary productivity and other ecosystem functions (Liu *et al*. 2018).

Despite overall trends in responses to environmental changes, some studies (including ours) find variable responses to warming (see Henry & Molau 1997, Erschbamer 2007, Elmendorf *et al*. 2012). A global review found that warming had positive effects on graminoid species abundance at cool sites but had neutral or negative effects at warm sites (Elmendorf *et al*. 2012). In our study, experimental warming influenced few communities, with only marginal support and limited effects on percent cover at the high elevation United States, suggesting that warming alone might be less influential to these communities, at least over short (*<*5 year) time scales. Our study also differs from some previous studies as we did not add ambient temperature of each year as a predictor or covariate in our model which may illuminate some effects of warming (Elmendorf *et al*. 2012). In addition to direct impacts on individuals, if the direction or magnitude of response to temperature varies among species, a change in temperature can also alter species interactions (Kordas *et al*. 2011). For example, in high elevation New Zealand, which showed support for the interaction treatment, if the ‘average’ species had 10% previous percent cover in the warming treatment this species would drop to 6.37%, and species in the removal treatment would slightly increase to 6.81% compared to ambient plots at 6.58%. In the combined warming and removal treatment at this site, the same ‘average’ species would have 5.35% showing a more pronounced decrease in percent cover when both factors are combined. This was the only site in our study which showed support (through model comparison) for the Warming × Removal treatment suggesting that, while each disturbance alone had impacts on some plant communities, they rarely seem to interact. This runs contrary to both our original expectation of synergistic effects and to some previous studies (Shevtsova & Ojala 1997, Virtanen *et al*. 2010). Nonetheless, this pattern of community composition modulating the effects of climate change is not consistent across experiments from different regions (Hobbie *et al*. 1999). Clearly, the response of alpine-plant communities and the role biotic and abiotic factors play in shaping those communities, is context-dependent.

As we saw here, community-wide responses suggest one trend while species within these communities respond variably to biotic and abiotic drivers. We found that the loss of the dominant species and experimental warming in these alpine-plant communities can have a variety of effects on the remaining community depending on species identity and location. While this result may not be novel to those familiar with the system (Pauli *et al*. 2014, Erschbamer 2007), our results can help reconcile previous conclusions by demonstrating that alpine-plant communities may broadly differ in the drivers structuring their communities. A pressing future challenge will be to understand the biological reasons behind these patterns of responses to abiotic and biotic disturbances, the combination of the two, or none of the above. We suggest that future studies focus on this overlapping role of biotic and abiotic disturbances in similar communities to determine if and how this pattern is affected by community composition. We also point to short- and long-term responses as an avenue for future research to explore how these patterns may be affected by the time-scale of the study. Previous warming experiments have drawn variable conclusions that are difficult to synthesize since studies often differ in the length of time or have varying responses throughout the length of a single study (Elmendorf *et al*. 2012, Kremers *et al*. 2015, Chapin III *et al*. 1995). Some previous experiments showed delayed response of subdominant groups to functional group removal treatment up to five years, with onset of biomass compensation depending on functional groups removed and remaining (McLaren & Turkington 2011). There are multiple mechanisms for time lags in the response of alpine-plants to climate change including establishment lag, extinction lag, and dispersal lag, all of which are affected by individual species physiology, demography, biotic interactions, and the physical environment (Alexander *et al*. 2018, Nomoto & Alexander 2021). As long-term data sets, such as this one, continue to grow, they offer opportunities for fruitful research ventures into different temporal and observational scales (species, community, and functional-group levels). Though a major conclusion of ‘it depends’ might feel unsatisfactory, in reality, it points to the complicated and multifaceted determinants of dynamics in diverse ecological communities.

## Acknowledgments

We would like to thank those who completed the field data collection for this project, specifically Daniela Aguirre, Kenna Rewcastle, Jeremiah Henning, Raina Fitzpatrick. The Stouffer Lab and many others who provided feedback on early stages of this project and manuscript. This project was made possible by a grant from the Marsden Fund Council from New Zealand Government funding, which is managed by the Royal Society Te Apa̅rangi (16-UOC-008 awarded to DBS), a University of Canterbury Doctoral Scholarship awarded to MLM, and University of Texas at El Paso Start-up funds to JRM. Fieldwork and larger project funding includes the Carlsberg Foundation and the Swedish FORMAS program. We thank Peter le Roux and two other anonymous reviewers who provided instrumental feedback on a previous version of this manuscript.

