## Supplemental for "Variable responses of alpine-plant communities to warming and loss of dominant species"

July 21, 2024

#### S1 Methods

Data used in this study form part of a larger study of alpine plant communities that combines experimental warming and dominant plant species removal among multiple globally-distributed elevation gradients (WaRM: Warming and (species) Removal in Mountains)[1]. Non-vascular plants such as mosses were included in the analysis. Mosses were only observed in low elevation CH plots and accounted for an average cover of 7.57%.

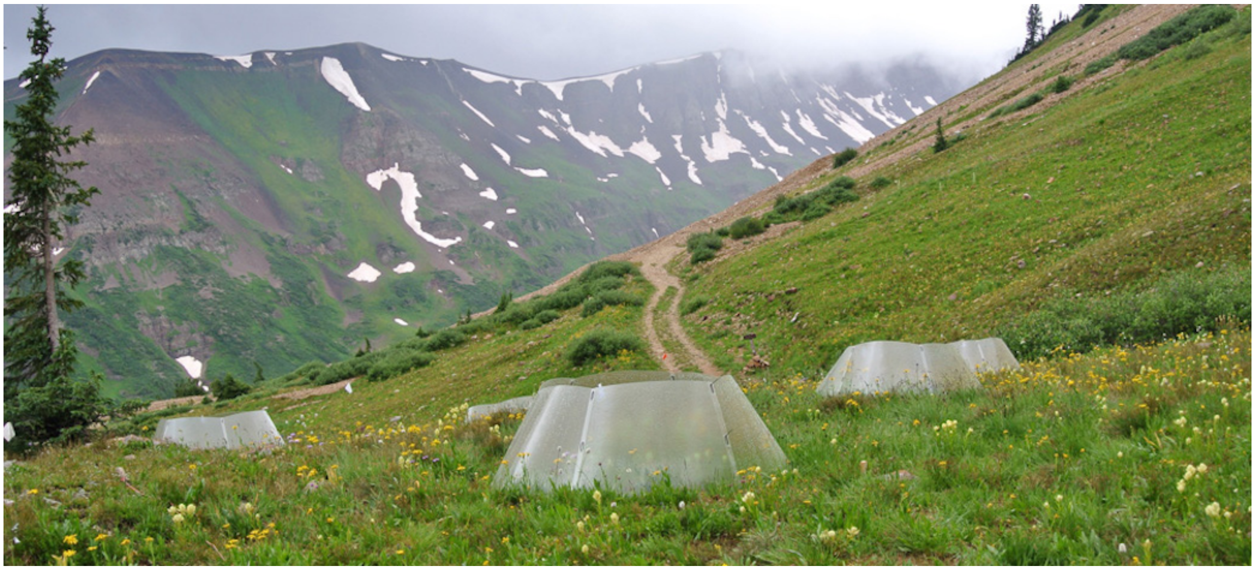

Fig. S1: View of the Colorado, US field site. Shows open-top chambers in place over experimental plots.

Table S1: MetaData for sites abbreviated using their two letter ISO code. Number of species records the number of species investigated in the model. Approximate length of the growing season. Average percent cover of a plot at the site including all species originally present. Amount modelled shows the average community percent cover in a plot actually included in the model after exclusions (i.e. those that were considered widespread species by our criteria) described in the methods.

| Location | Elevation (meters) | No. of Plots | Latitude | Longitude | Growing season (months) | Number of species | Average Percent Cover | Percent Cover Modelled |
|---|---|---|---|---|---|---|---|---|
| CA | Low (1431) | 32 | 60.979 | -138.408 | 3 | 16 | 82.9 | 64.1 |
| | High (1900) | 32 | 60.954 | -138.423 | 3 | 7 | 59.7 | 38.4 |
| CH | Low (2101) | 40 | 46.77497 | 9.862969 | 4 | 12 | 126.0 | 83.0 |
| | High (2353) | 40 | 37.707 | 101.372 | 4 | 12 | 108.0 | 78.5 |
| CN | Low (3200) | 32 | 37.617 | 101.200 | 4 | 19 | 127.0 | 89.5 |
| | High (4004) | 32 | 46.774113 | 9.856959 | 4 | 11 | 146.0 | 81.8 |
| NZ | Low (1071) | 32 | -39.296 | 175.727 | 6 | 26 | 106.0 | 85.9 |
| | High (1611) | 32 | -39.285 | 175.623 | 6 | 9 | 29. 3 | 12.8 |
| US | Low (2740) | 40 | 38.715 | -106.823 | 3 | 18 | 88.7 | 53.8 |
| | High (3460) | 40 | 38.992 | -107.067 | 3 | 18 | 96.6 | 73.9 |

#### S1.1 Dominant Species

The dominant species at each site was not included in analyses for three main reasons. First, it is statistically complicated to include this as the model estimates the individual parameters for each species under each treatment but the dominant is inherently absent from two of the treatments. Second, by removing the dominant we lose little information about the remaining plots since they account for between 10.2 and 52.3, with a median of 20 percent cover in non-removal plots at each sites. While these may seem like large numbers, total cover in plots ranged from 33.5 to 121.7 percent cover.

Table S2: Average percent cover of the dominant species per plot at each site compared to cover of the total community. Shows average percent cover in each treatment (across all plots and years studied), as well as its average percent cover before exclusion in the removal and removal:warming treatment. These values are an average of all years and are expected to be low in removal treatments due to years of removal of adults. Total Community shows the average percent cover of a plot summed across of all species (including the dominant species).

| Location | Elevation | Species | Ambient | Warming | Removal | Removal:Warming | Total Community |
|---|---|---|---|---|---|---|---|
| CA | Low | Salix reticulata | 19.87 | 14.4 | 7.06 | 6.69 | 88.48 |
| | High | Carex consimilis | 28.0 | 30.5 | 10.5 | 9.47 | 58.91 |
| CH | Low | Vaccinium uliginosum | 33.2 | 52.3 | 16 | 18.6 | 121.7 |
| | High | Vaccinium uliginosum | 34.1 | 35.6 | 15.3 | 12.5 | 111.87 |
| CN | Low | Stipa aliena | 11.6 | 14.6 | 12.6 | 12.3 | 109.10 |
| | High | Kobresia pygmaea | 10.2 | 12.1 | 12.1 | 6.95 | 113.49 |
| NZ | Low | Calluna vulgaris | 18.1 | 23.5 | 7.83 | 7.88 | 108.77 |
| | High | Gaultheria collensoi | 29.1 | 21.8 | 6.43 | 4.78 | 33.5 |
| US | Low | Wyethia amplexicaulis | 20.2 | 37.87 | 5.03 | 4.08 | 80.94 |
| | High | Juncus drummondii | 21.6 | 16.07 | 3.22 | 2.33 | 87.97 |

Third, analysis of these species in non-removal treatments at their respective sites revealed that their dynamics do not vary as a function of the warming treatment; therefore, their inclusion in the analysis would not alter the overall conclusions about community dynamics. In two of the ten sites, the warming treatment influenced the dominant: in the US (high elevation) and CH (low elevation), based on the same model weighting procedure as in the main text (see Methods: Model Comparison). Despite this support for warming treatment in the best-fit model, warming the dominant species showed small changes in growth as a response. For example, at low elevation CH density-dependent change in cover increased from $\exp(g|A) = 0.42$ under Ambient conditions to $\exp(g|W) = 0.52$ under warming conditions.

#### S1.2 Wide-spread species

As noted in the Methods, we also concentrated our analysis on the most widespread species within each site. We defined widespread as the species observed in at least 25 percent of total plots (for example: $0.25 \times 32$ plots $\times$ 4 years = 32 plots, which becomes the minimum number of plots a species must be present in to be analyzed). The resulting number of plots varied by location depending on the number of years studied (Supplemental Table S1). We tested the robustness of this threshold, and found that fitting the model to species in 20 and 30% of plots showed qualitatively consistent results at the site level (results not shown). Widespread species represent those that tend to be pervasive and abundant. While these widespread species represent a subset of the total species observed, they account for the majority of the percent cover observed in the plots (Supplemental Table S1 Supplemental Fig S2 and S3).

#### S1.3 Model Inference

Our Bayesian, two-level hierarchical model for our dynamic model of year-to-year variation in cover of a focal species $i$ within a sampled plot may be written as:

$$N_{i,t} \sim \text{Beta}(\lambda_{i,t}, \phi, z_i) \tag{S1}$$

$$\lambda_{i,t} \sim Q_i + N_{i,t-1} e^{G_{i|R,W}} \tag{S2}$$

$$Q_i \sim \frac{0.5}{1 + e^{-(q_{common} + \Delta q_i)}} \tag{S3}$$

$$\{q_{common}, g, g_R, g_W, g_{RW}\} \sim \text{Normal}(0, 10) \tag{S4}$$

$$\{\Delta q_i, \Delta g_i, \Delta g_{R_i}, \Delta g_{W_i}, \Delta g_{RW_i}\} \sim \text{Multivariate Normal}(\sigma, \rho) \tag{S5}$$

$$\sigma \sim \text{HalfCauchy}(0, 2) \tag{S6}$$

$$\rho \sim \text{LKJcorr}(2) \tag{S7}$$

$$\phi \sim \text{Gamma}(0.01, 0.01) \tag{S8}$$

$$z_i \sim \text{Beta}(1, 1) \tag{S9}$$

where $\lambda_{i,t}$ refers to the mean predicted percent cover of species $i$ in year $t$.

We fit our models using a zero-inflated beta distribution. A beta distribution is appropriate for percent cover data as it is ideal for continuous data and allows for a flexible shape (U-shaped, L-shaped, etc.)[2,3] (Eqn S1). Here our weakly informative priors help the Markov chains' convergence and stabilization while still letting the data speak by excluding various 'unphysical' possibilities that would otherwise take over the posterior distribution[4]. Rather than estimate each species' deviations from the grand mean ($\Delta$ parameters) as separate fixed effects, we parameterized these deviations in a comparable manner to how random effects are included in mixed-effects models.

We used weakly informative priors to parametrize the treatment (control, R, W, and RW) terms within growth (Eqn S5) since we had no previous knowledge with which to constrain these parameters. We also defined varying effects of each focal species, that correspond to deviations from the grand mean ($\Delta$ parameters), with a non-centered parametrization of a multivariate normal distribution[5] with a mean ($\sigma$) and covariance matrix ($\rho$) between the varying effects for each species (Eqn S6). We used weakly informative priors of half Cauchy for the mean and LKJcorr(1) for the covariance matrix (Eqn S7-S8)[6]. The Cauchy distribution is a thick-tailed probability distribution; in this case, a half-Cauchy (restricted to positive values) acts as a weakly informative/regularizing prior for standard deviations[6]. The LJKcorr distribution of correlation coefficients provides a weakly informative prior on the covariance matrix which is skeptical of extreme correlations near -1 or 1[7]. This implies that our model is skeptical of extreme correlations between the species since we wish to allow each species to respond to treatments individually.

### S2 Results

#### S2.1 Community level results

Models utilized here performed well across a range of observations, species richness (7–26), and plant species across sites (Low Main Text Fig 1 and High Fig S5 ).

In high elevation CN, the Removal × Warming model received 20% WAIC weight; however, this site showed mixed support for all treatment models with them receiving an equal split of the weight possibly due to the low sample size at this site.

Density-independent influx also varied by site with NZ having the lowest average contribution to species cover and CN having the highest. Other sites received around 1% influx. Density-independent influx of species in these communities which may have resulted from an influx of seeds, germination of seeds from the seed bank, or lateral encroachment of neighboring plants. While a small amount, given density-dependent change in cover of less than 1 in some areas, influx may be allowing plots to maintain cover.

Table S3: Fixed effects of linear mixed effects models showing the relationship between change in percent cover and treatments. ANOVA showed no effect of treatment (F(3,251)=1.202, p=0.31)

| Factor | Value | Std. Error | DF | t-value | p-value |
|---|---|---|---|---|---|
| Control | 12.66 | 12.78 | 244 | 0.99 | 0.32 |
| Removal | 4.09 | 4.41 | 244 | 0.929 | 0.35 |
| Warming | -5.78 | 4.39 | 244 | -1.31 | 0.18 |
| Removal:Warming | 4.84 | 4.39 | 244 | 1.10 | 0.27 |

#### S2.2 Species-specific results

Species responded idiosyncratically to treatments and showed differing rates of influx of species. While in some sites responses of individual species to treatments mimicked the grand means (e.g. Fig S8), theses species still showed a range of changes in percent cover. In some sites an individual species showed a relatively large change in its percent cover in response to the removal of the dominant species (for example) even though the grand means at that site showed minimal responses to treatments (e.g. Fig S6). These results illustrate that response of the whole community does not dictate the response of individual species. Figures presented for each site below. Sites with support for multiple models have are depicted with averaged posteriors predictions. If a treatment was not supported in the best-fit model(s) that treatment represents the ambient effect if a single treatment or the single treatment if the combination effect.

#### S2.3 Linear Regression Analysis

We examined the change in cover as the total cover of a plot (the sum of all species' percent cover) as final (last year measured) minus initial (first year measured). China is not represented as many plots in this location were not recorded in the first and last year leading to a low and unbalanced sample. We used an ANOVA to determine if there were any differences among treatment across all locations and found no effects of treatment (ANOVA, F(3,251)=1.202, p=0.31, Fig. S4). To examine the effects treatments within sites, we used a linear mixed effects model with elevation nested within location as a random effect. This test did not reveal any significant effect of treatment (Table S3).

Total percent cover in experimental plots remained similar throughout our study and did not vary with treatment (Fig S2). In Canada (CA) and Switzerland (CH), most plots showed no difference between final and initial total percent cover at the community-level. United States (US) plots increased in percent cover, and New Zealand (NZ) plots showed variable results (Fig S2). While percent cover in plots varied across locations and elevations, treatment was not a significant predictor (Table S3). China (CN) was not used in this analysis as it had too few replicates of the same plot measured in both the final and initial years. For more details see Supplemental (S2.3 Linear Regression Analysis).

#### S2.4 Functional Group Analysis

To examine how treatments affected groups of plants rather than individual species, we lumped plants into their functional form: C3 graminoid, forb, legume, and woody shrub based on taxonomic identifications. Similarly to focal-species analysis preformed in the main text, we used a population-dynamics model for percent cover of a functional group as a function of previous percent cover. Here our model format allows for varying effects of each functional group that correspond to deviations from the grand mean ($\Delta$ parameters). Unlike species specific analysis, since there are inherently fewer functional groups we were able to combine data across sites and analyze the effect of site (elevation within location) as a random factor with ten levels. Again, we used a Beta distribution to generate predicted cover since our observations are always a series of non-negative integers and weakly informative priors for all of the parameters. We modeled cover with our linear mathematical model of population dynamics (Eqn 1 and 2 in the main text), we used weakly informative priors to parametrize the treatment (control, R, W, and RW) terms within growth (Eqn 2 in the main text) since we had no previous knowledge with which to constrain these parameters. We performed sampling to determine the posterior distributions of model parameters through the function "brm" from the package "brms"[8] in the statistical program R (version 3.4.2)[9]. We ran two chains with a warm-up of 1000 iterations and 3000 sampling iterations each, and this produced a final combined posterior of 6000 MCMC samples for each model. We determined that parameters converged when trace plots were well mixed and stationary, and the Gelman-Rubin convergence diagnostic equaled one[10,11]. As with the species specific analysis, we compared a series of models (developed in a step-wise fashion from least parameterized to most) to examine the relative importance of each treatment and treatment combination to variation in population dynamics (see main text Table 1). Best-fit models were those with the lowest WAIC and an Akaike weight greater than 0.8.

This analysis focused on five functional groups (forb, C3 Graminoid, woody shrub, legume, and moss) as a random effect similar to the analysis in the main text where focal species was a random effect. Additionally this analysis included a random effect for site. Here the winning models was the null model (0.83 WAIC weight). Showing no support for any treatment at the functional group level. The recruitment models did not converge in this analysis.

Table S4: WAIC comparison of candidate models on functional group data. The Null model received the most support and warming was the only treatment to receive any support with 5% of the weight. These models are the same formulation as the main text Table 1 but here $i$ in $\Delta_i$ refers to a functional group. Model comparisons is also the same procedure as the main text (see Methods: Model Comparison section)

| Model Name | WAIC | pWAIC | Weight |
|---|---|---|---|
| Null | -1561.12 | 64.58 | 0.83 |
| Ambient | -1557.25 | 69.12 | 0.12 |
| Warming | -1555.47 | 72.87 | 0.05 |
| Removal | -1548.63 | 55.98 | 0 |
| Removal + Warming | -1547.86 | 61.19 | 0 |
| Removal × Warming | -700.43 | 4.99 | 0 |

Table S5: Species list by elevation shows species and functional group assignments for the species investigated at each site.

| Location | Elevation | Taxa | Functional Group |
|---|---|---|---|
| CA | Low | *Achillea millefolium* | Forb |
| | | *Aconitum delphinifolium* | Forb |
| | | *Anemone* spp. | Forb |
| | | *Artemesia norvegica* | Forb |
| | | *Carex* spp. | C3 Graminoid |
| | | *Castilleja* spp. | Forb |
| | | *Festuca altaica* | C3 Graminoid |
| | | *Lupinus arcticus* | Legume |
| | | *Mertensia paniculata* | Forb |
| | | *Myosotis alpestris* | Forb |
| | | *Polemonium pulcherrimum* | Forb |
| | | *Senecio lugens* | Forb |
| | | *Valeriana capitata* | Forb |
| | | *Veronica wormskjoldii* | Forb |
| | High | *Equisetum variegatum* | Forb |
| | | *Pedicularis* spp. | Forb |
| | | *Petasites frigidus* | Forb |
| | | *Salix reticulata* | Woody |
| | | *Salix rotundifolia* | Woody |
| | | *Saxifraga oppositifolia* | Forb |
| | | *Senecio atropurpureus* | Forb |
| CH | Low | *Avenella flexuosa* | C3 Graminoid |
| | | *Calamagrostis villosa* | C3 Graminoid |
| | | *Homogyne alpina* | Forb |
| | | *Luzula sieberi* | C3 Graminoid |
| | | *Oxalis acetosella* | Forb |
| | | *Empetrum nigrum* subsp. *hermaphroditum* | Woody |
| | | *Rhododendron ferrugineum* | Woody |

| | | | |
|---|---|---|---|
| | | *Rumex alpestris* | Forb |
| | | *Vaccinium myrtillus* | Woody |
| | | *Vaccinium vitis-idaea* | Woody |
| | High | *Anthoxanthum alpinum* | C3 Graminoid |
| | | *Arnica montana* | Forb |
| | | *Carex curvula* | C3 Graminoid |
| | | *Diphasiastrum alpinum* | Moss |
| | | *Empetrum nigrum* subsp. *hermaphroditum* | Woody |
| | | *Gentiana punctata* | Forb |
| | | *Helictotrichon versicolor* | C3 Graminoid |
| | | *Hieracium alpinum* | Forb |
| | | *Homogyne alpina* | Forb |
| | | *Leontodon helveticus* | Forb |
| | | *Ligusticum mutellina* | Forb |
| | | *Loiseleuria procumbens* | Woody |
| | | *Luzula lutea* | C3 Graminoid |
| | | *Nardus stricta* | C3 Graminoid |
| | | *Phyteuma hemisphaericum* | Forb |
| | | *Senecio incanus* | Forb |
| | | *Vaccinium myrtillus* | Woody |
| CN | Low | *Ajania tenuifolia* | Forb |
| | | *Aster flaccidus* | Forb |
| | | *Elymus nutans* | C3 Graminoid |
| | | *Euphrasia regelii* | Forb |
| | | *Gentiana aristata* | Forb |
| | | *Gentiana straminea* | Forb |
| | | *Gueldenstaedtia diversifolia* | Legume |
| | | *Kobresia humilis* | C3 Graminoid |
| | | *Lancea tibetica* | Forb |
| | | *Leontopodium nanum* | Forb |
| | | *Morina chinensis* | Forb |
| | | *Oxytropis qinghaiensis* | Forb |

| | | | |
|---|---|---|---|
| | | *Poa crymophila* | Forb |
| | | *Potentilla bifurca* | Forb |
| | | *Potentilla saundersiana* | Forb |
| | | *Saussurea nigrescens* | Forb |
| | | *Saussurea superba* | Forb |
| | | *Stellaria umbellata* | Forb |
| | | *Taraxacum mongolicum* | Forb |
| | | *Thalictrum alpinum* | Forb |
| | | *Thalictrum rutifolium* | Forb |
| | High | *Allium sikkimense* | Forb |
| | | *Anaphalis lactea* | Forb |
| | | *Aster flaccidus* | Forb |
| | | *Carex przewalskii* | C3 Graminoid |
| | | *Kobresia humilis* | C3 Graminoid |
| | | *Lancea tibetica* | Forb |
| | | *Leontopodium nanum* | Forb |
| | | *Oxytropis qinghaiensis* | Legume |
| | | *Pedicularis kansuensis* | Forb |
| | | *Poa crymophila* | C3 Graminoid |
| | | *Poa orinosa* | C3 Graminoid |
| | | *Potentilla saundersiana* | Forb |
| | | *Rheum pumilum* | Forb |
| US | Low | *Achillea millefolium* | Forb |
| | | *Alopecurus pratensis* | C3 Graminoid |
| | | *Elymus elymoides* | C3 Graminoid |
| | | *Eremogone congesta* | Forb |
| | | *Erigeron* sp. | Forb |
| | | *Erigeron speciosus* | Forb |
| | | *Festuca thurberi* | C3 Graminoid |
| | | *Galium septentrionale* | Forb |
| | | *Poa fendleriana* | C3 Graminoid |
| | | *Potentilla gracilis* | Forb |

| | | | |
|---|---|---|---|
| | | *Rosa woodsii* | Woody |
| | | *Taraxicum officinale* | Forb |
| | | *Tragopogon dubius* | Forb |
| | | *Vicia americana* | Legume |
| | High | *Agoseris glauca* | Forb |
| | | *Arctostaphylos uva-ursi* | Woody |
| | | *Arnica mollis* | Forb |
| | | *Carex ebenea* | C3 Graminoid |
| | | *Castilleja sulphurea* | Forb |
| | | *Draba spectabilis* | Forb |
| | | *Erigeron glacialis* | Forb |
| | | *Erythronium grandiflorum* | Forb |
| | | *Poa alpina* | C3 Graminoid |
| | | *Poa arctica* | C3 Graminoid |
| | | *Senecio crassulus* | Forb |
| | | *Sibbaldia procumbens* | Forb |
| | | *Viola labradorica* | Forb |
| NZ | Low | *Asteraceae* | Forb |
| | | *Celmisia glandulosa* | Forb |
| | | *Celmisia gracilenta* | Forb |
| | | *Celmisia spectabilis* | Forb |
| | | *Chionochloa pallens* | C3 Graminoid |
| | | *Chionochloa rubra* | C3 Graminoid |
| | | *Coprosma cheesemanii* | Woody |
| | | *Coprosma perpusilla* | Woody |
| | | *Dracophyllum recurvum* | Woody |
| | | *Dracophyllum subulatum* | Woody |
| | | *Epacris alpina* | Woody |
| | | *Euphrasia cuneata* | Forb |
| | | *Gaultheria colensoi* | Woody |
| | | *Gonocarpus micranthus* | Forb |
| | | *Veronica tetragona* subsp. *subsimilis* | Woody |

| | | |
|---|---|---|
| | *Veronica venustula* | Woody |
| | *Leucopogon fraseri* | Woody |
| | Orchids | Forb |
| | *Oreobolus strictus* | Forb |
| | *Pentachondra pumila* | Woody |
| | *Poa colensoi* | C3 Graminoid |
| | *Wahlebergia pygmaea* | Forb |
| High | *Anistome aromatica* | Forb |
| | *Celmisia gracilenta* | Forb |
| | *Chionochloa pallens* | C3 Graminoid |
| | *Gentianella bellidifolia* | Forb |
| | *Luzula colensoi* | Forb |
| | *Muehlenbeckia axillaris* | Woody |
| | *Poa colensoi* | C3 Graminoid |
| | *Raoulia albosericea* | Forb |
| | *Wahlebergia pygmaea* | Forb |

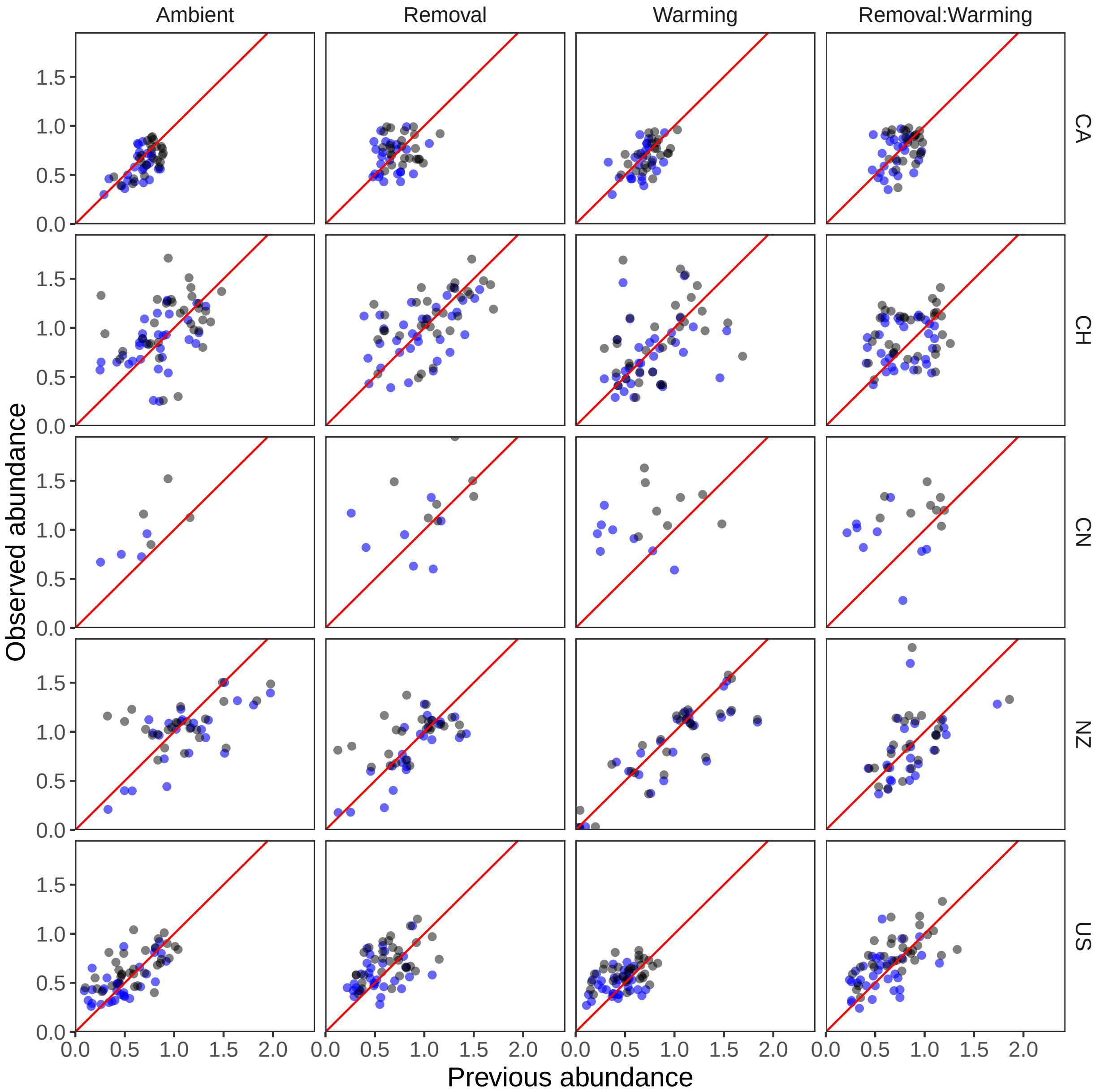


Fig. S2: Raw and modeled data on a plot level at each site at the low elevation. Shows total percent cover in a plot as gray dots and the percent cover used in the analysis (after removal of the dominant species and less widespread species) as blue dots. The red line represents 1:1 or where previous cover would equal current cover i.e. when plots neither grow nor decline in percent cover. Data scaled for beta distribution (i.e. between 0-1 for each species).

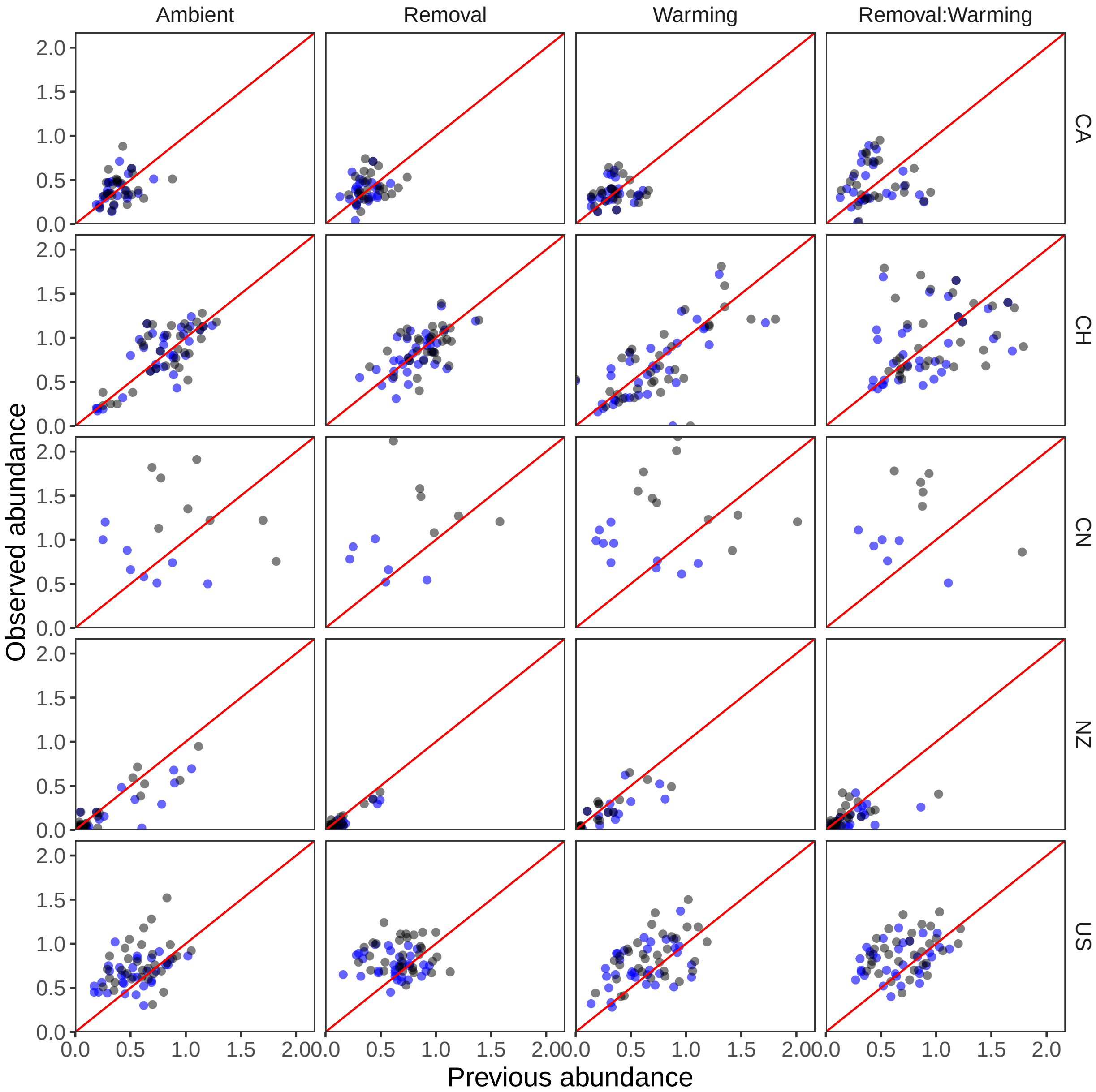


Fig. S3: Raw and modeled data on a plot level at each site at the high elevation. Shows total percent cover in a plot as gray dots and the percent cover used in the analysis (after removal of the dominant species and less widespread species) as blue dots. The red line represents 1:1 or where previous cover would equal current cover i.e. when plots neither grow nor decline in percent cover. Data scaled for beta distribution (i.e. between 0-1 for each species).

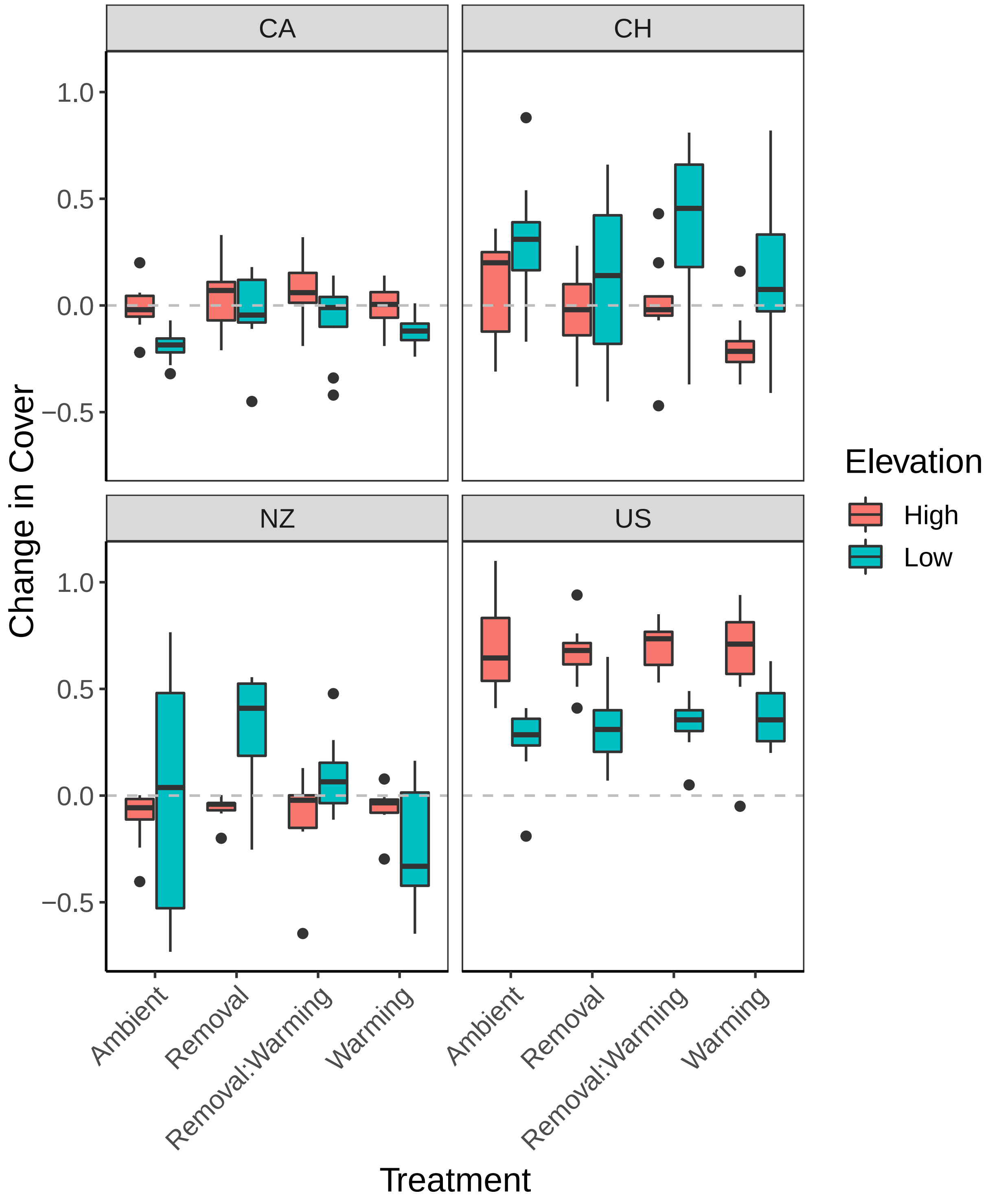


Fig. S4: Change in percent cover within plots across treatments and sites. Change is calculated as final (last year measured) minus initial (first year measured). Percent cover of all species is summed to reveal one value for each plot. China is not represented as many plots in this location were not recorded in the first and last year.

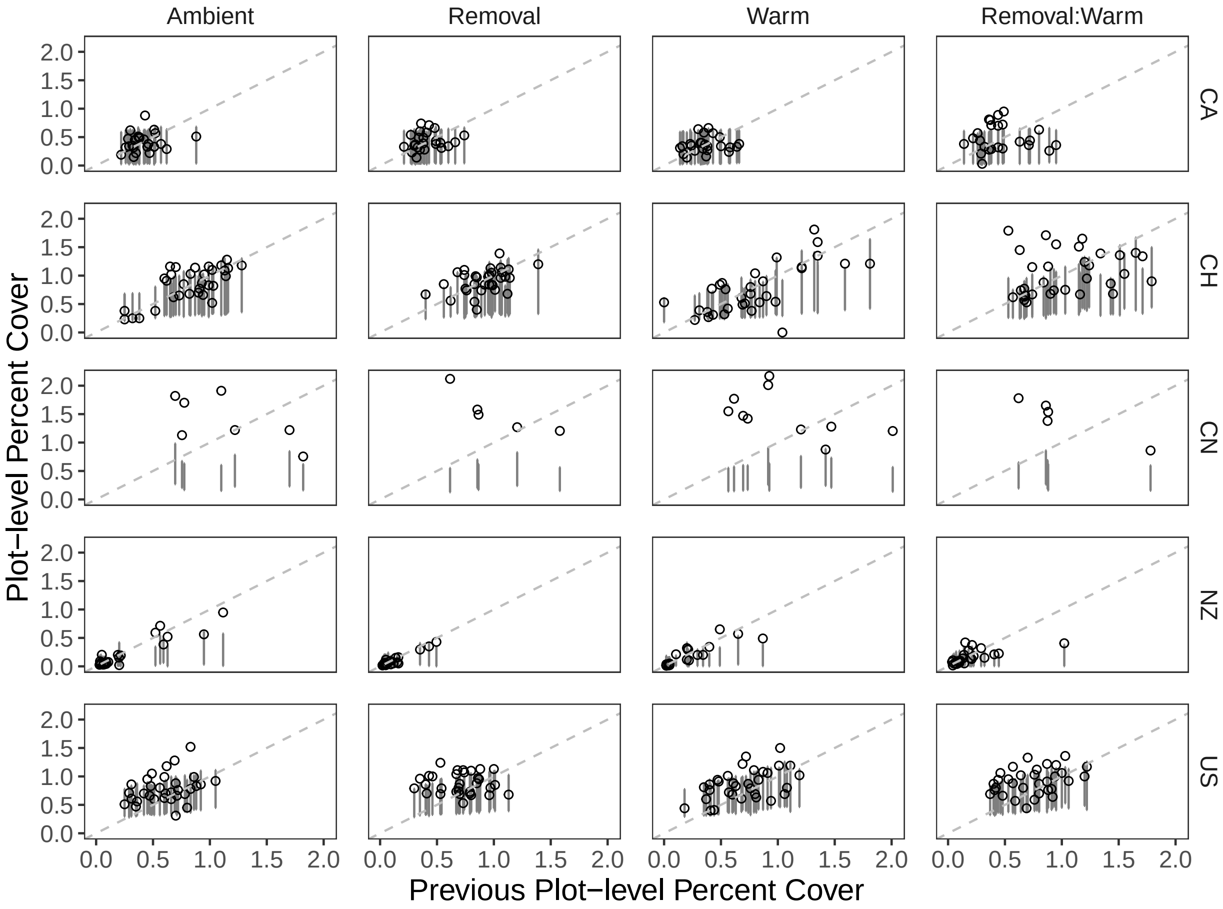


Fig. S5: Predicted and observed cover for all plots in high elevation across sites. Shows the model fit by overlaying predictions on top of observations. Points show the total observed cover in a plot, while shaded bars shows confidence interval (89%) of the predicted total cover for that plot. Predicted cover is calculated according to Eqn 1 then summed for each species within a plot to yield a plot level predicted cover. The dashed line represents 1:1 or where previous cover would equal current cover i.e. when plots neither grow nor decline in percent cover. Data scaled for beta distribution (i.e. between 0-1 for each species).

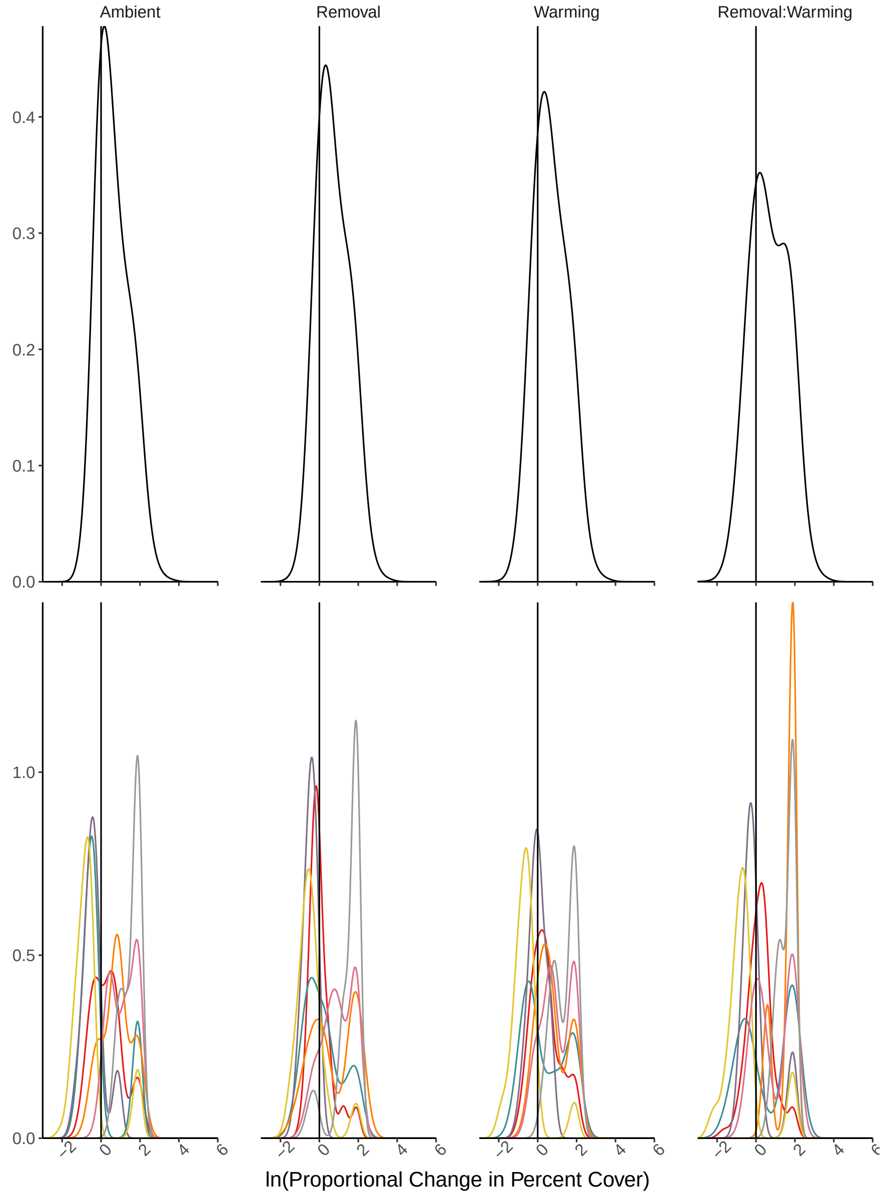


Fig. S6: Probability density of proportional change in percent cover within the high elevation Canada site. Logarithmic proportional change in cover is calculated as $(\ln(N_{i,t}/N_{i,t-1}) = \ln(Q_i/N_{i,t-1} + e^{G_{i|RW}}))$ as sampled from the joint posterior for parameters in our statistical model. A logarithmic change of zero ($\ln(1) = 0$; solid vertical line) represents no change over time suggesting that populations are at/near their equilibrium. The top row shows the predictions based on the community-level grand mean or the 'average' species, which in this case only shows minor effects of the removal treatment. The second row shows the predicted change in percent cover for all species at this site illustrating how species vary within and among treatments. This site has support for multiple models and the posterior predictions were averaged across winning models: Ambient, Warming, and Removal. Since this metric includes observed previous cover, differences seen in unsupported treatments reflect variations in species' cover rather than inferred parameters.

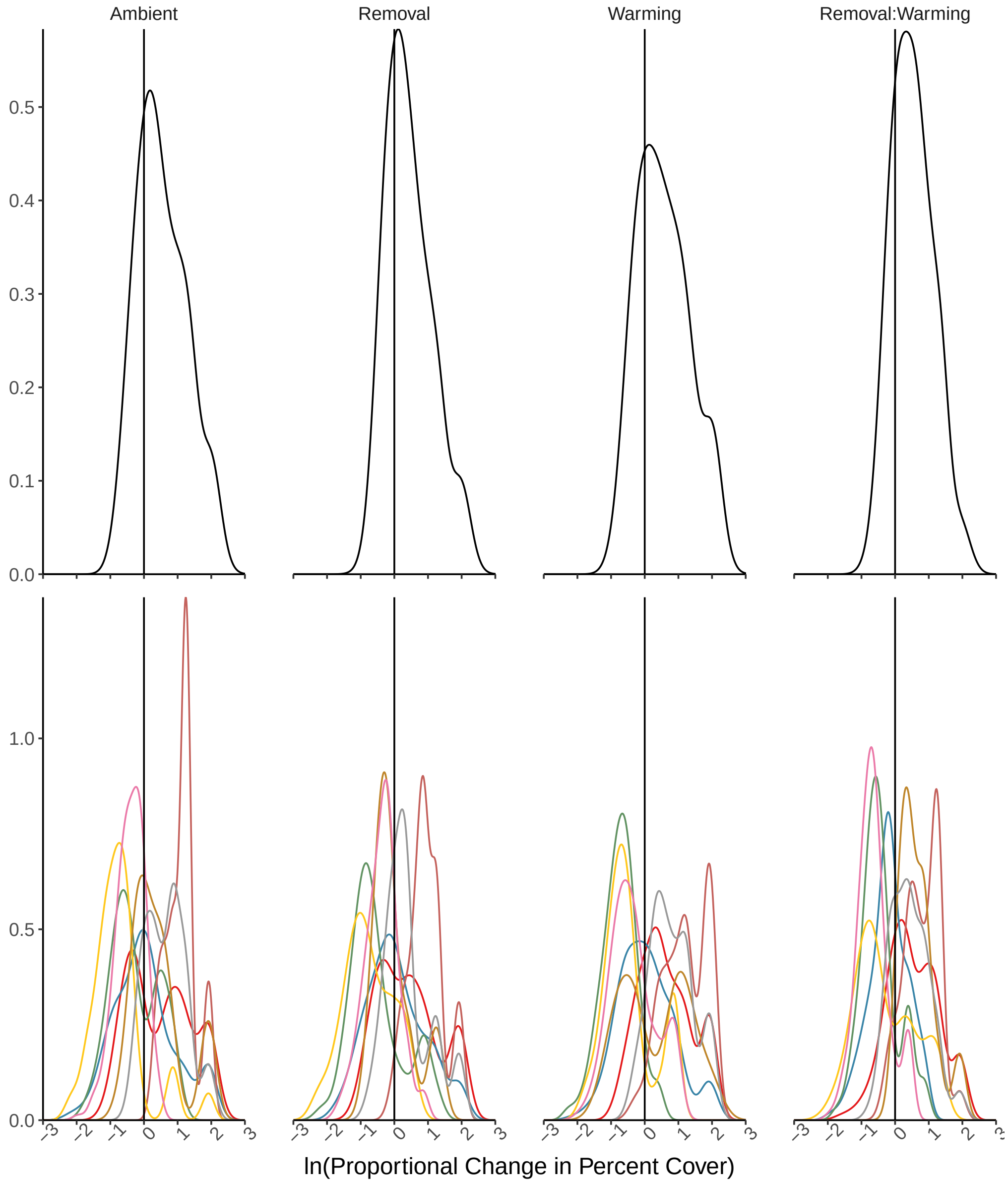


Fig. S7: Distribution of predicted change in percent cover within the low elevation Swiss site. Logarithmic proportional change in cover is calculated as $(\ln(N_{i,t}/N_{i,t-1}) = \ln(Q_i/N_{i,t-1} + e^{G_{i|RW}}))$ as sampled from the joint posterior for parameters in our statistical model. A logarithmic change of zero ($\ln(1) = 0$; solid vertical line) represents no change over time suggesting that populations are at/near their equilibrium. The top row shows the predictions based on the community-level grand mean or the 'average' species, which in this case only shows minor effects of the treatments. The second row shows the predicted change in percent cover for all species at this site illustrating how species vary within and among treatments. This site has support for multiple models and the posterior predictions were averaged across winning models: Removal + Warming, Removal, Removal × Warming, and Ambient. Since this metric includes observed previous cover, differences seen in unsupported treatments reflect variations in species' cover rather than inferred parameters.

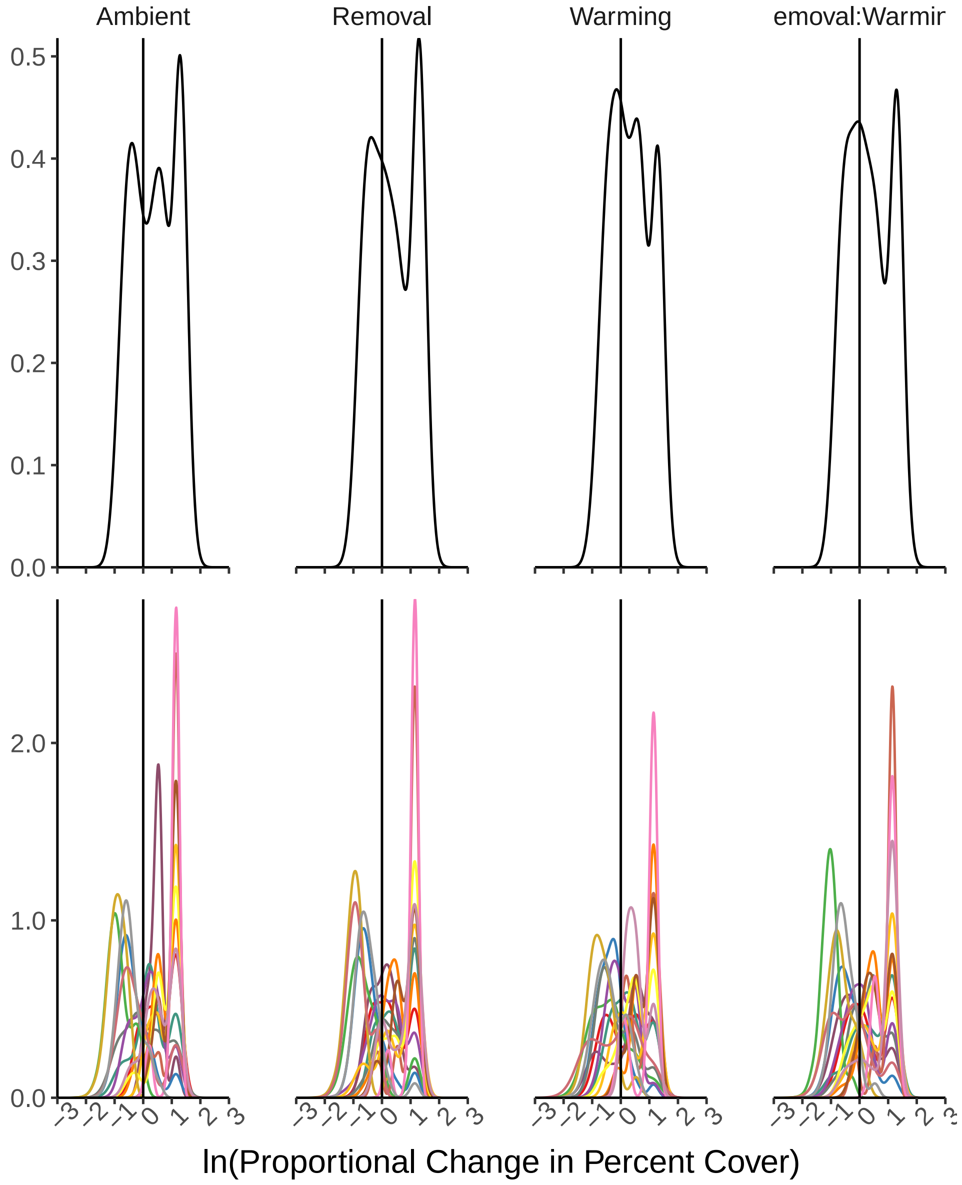


Fig. S8: Distribution of predicted change in percent cover within the high elevation Swiss site. Logarithmic proportional change in cover is calculated as $(\ln(N_{i,t}/N_{i,t-1}) = \ln(Q_i/N_{i,t-1} + e^{G_{i|RW}}))$ as sampled from the joint posterior for parameters in our statistical model. A logarithmic change of zero ($\ln(1) = 0$; solid vertical line) represents no change over time suggesting that populations are at/near their equilibrium. The top row shows the predictions based on the community-level grand mean or the 'average' species. The second row shows the predicted change in percent cover for all species at this site illustrating how species vary within and among treatments. Treatment or combination models did not receive any model support at this site so their predictions reflect the ambient predictions. Since this metric includes observed previous cover, differences seen in unsupported treatments reflect variations in species' cover rather than inferred parameters.

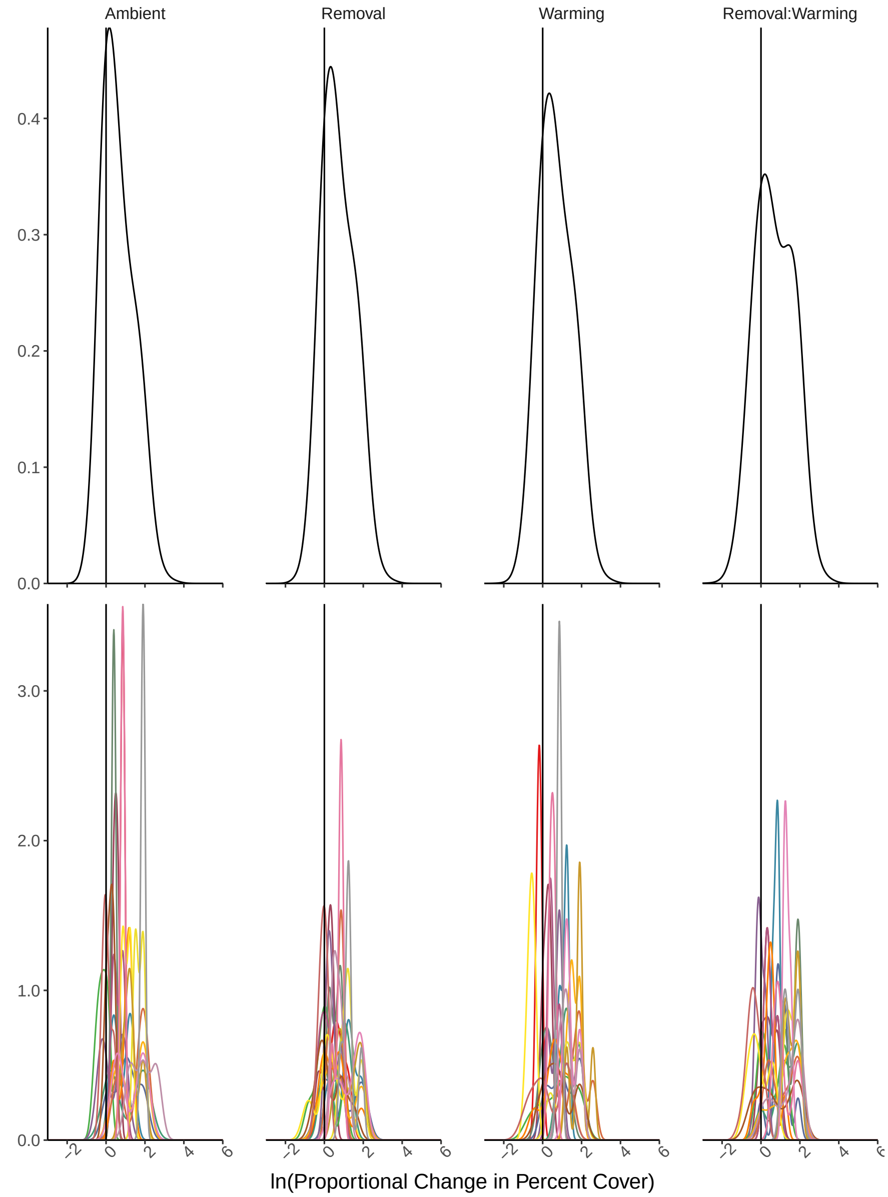


Fig. S9: Distribution of predicted change in percent cover within the low elevation China site. Logarithmic proportional change in cover is calculated as $(\ln(N_{i,t}/N_{i,t-1}) = \ln(Q_i/N_{i,t-1} + e^{G_{i|RW}}))$ as sampled from the joint posterior for parameters in our statistical model. A logarithmic change of zero ($\ln(1) = 0$; solid vertical line) represents no change over time suggesting that populations are at/near their equilibrium. The top row shows the predictions based on the community-level grand mean or the 'average' species. The second row shows the predicted change in percent cover for all species at this site illustrating how species vary within and among treatments. This site showed support for multiple models and the posterior predictions were averaged across winning models: Ambient, Removal, and Warming. Since this metric includes observed previous cover, differences seen in unsupported treatments reflect variations in species' cover rather than inferred parameters.

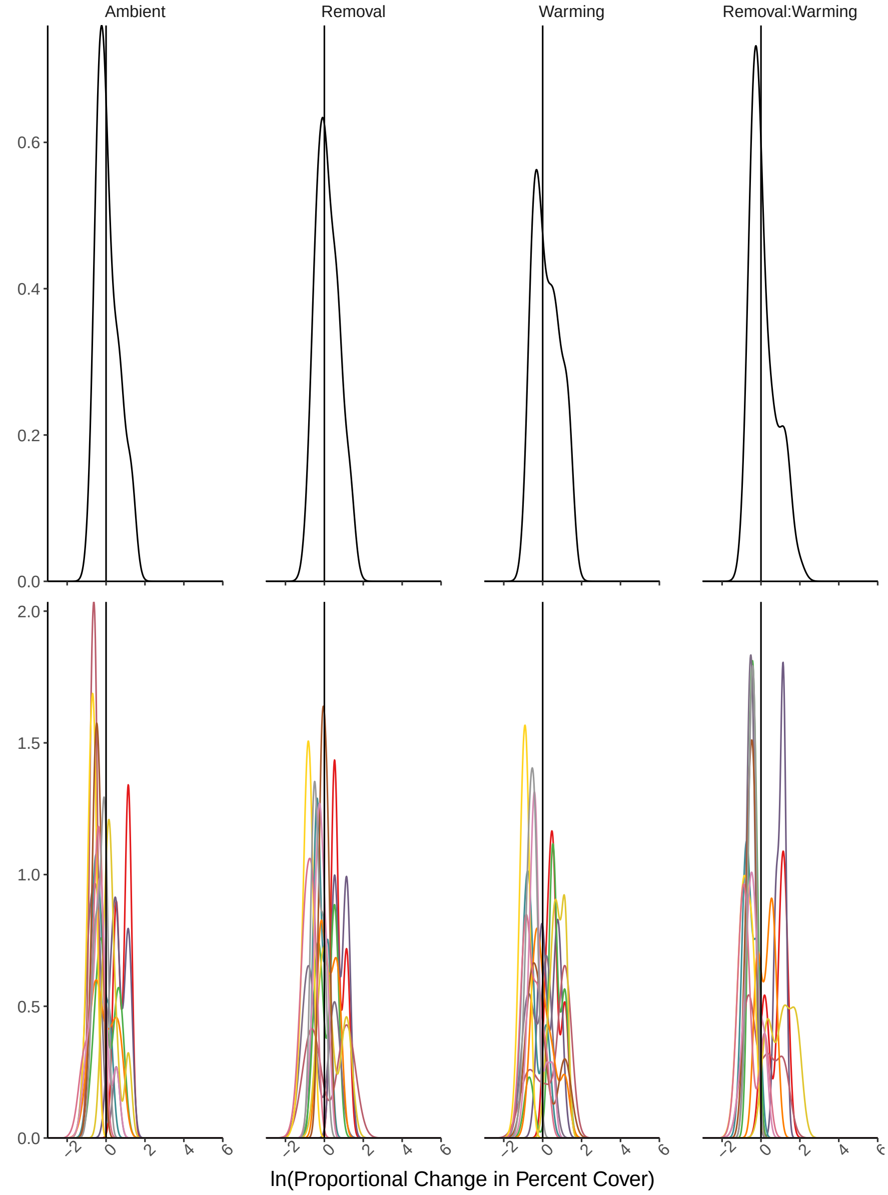


Fig. S10: Distribution of predicted change in percent cover within the low elevation New Zealand site. Logarithmic proportional change in cover is calculated as $(\ln(N_{i,t}/N_{i,t-1}) = \ln(Q_i/N_{i,t-1} + e^{G_{i|RW}}))$ as sampled from the joint posterior for parameters in our statistical model. A logarithmic change of zero ($\ln(1) = 0$; solid vertical line) represents no change over time suggesting that populations are at/near their equilibrium. The top row shows the predictions based on the community-level grand mean or the 'average' species. The second row shows the predicted change in percent cover for all species at this site illustrating how species vary within and among treatments. This site showed some support for all treatment and interaction models and this figure reflects averaged parameter estimates.

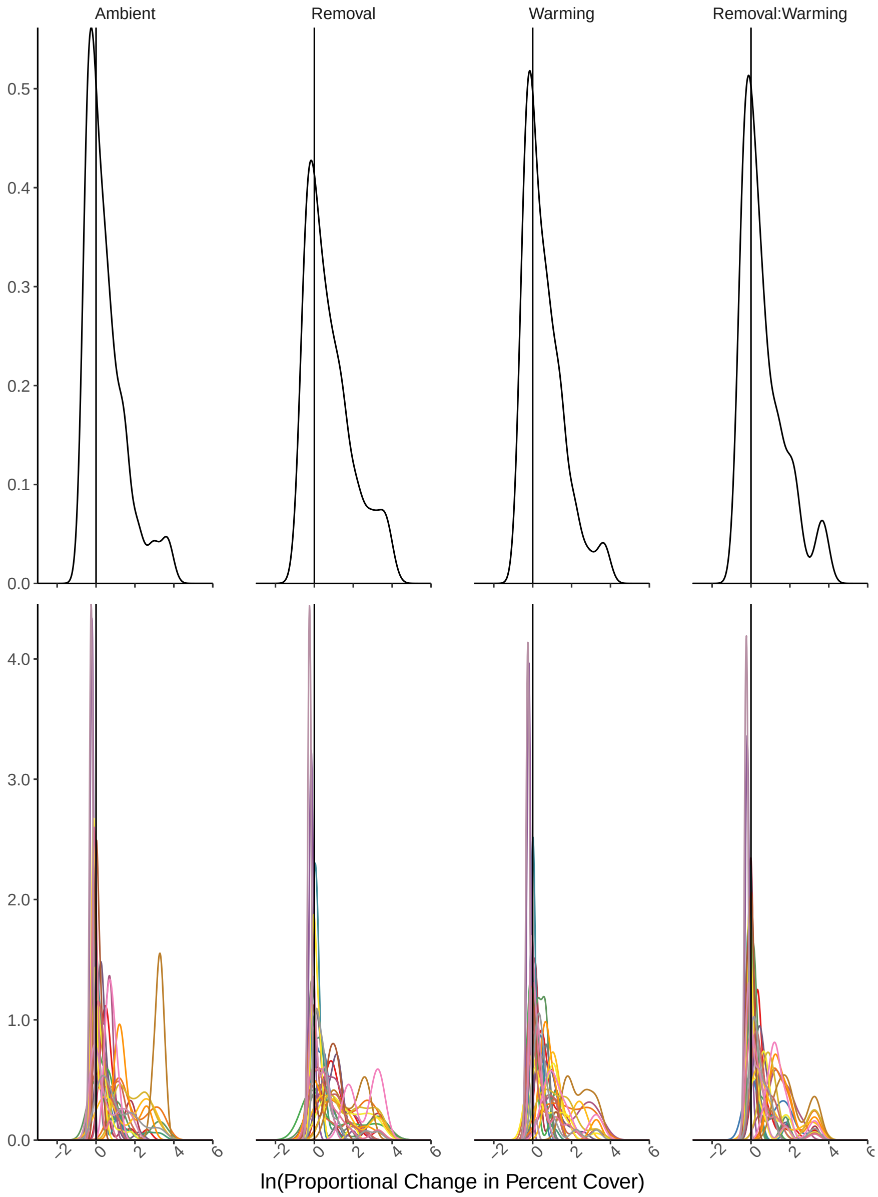


Fig. S11: Distribution of predicted change in percent cover within the low elevation New Zealand site. Logarithmic proportional change in cover is calculated as $(\ln(N_{i,t}/N_{i,t-1}) = \ln(Q_i/N_{i,t-1} + e^{G_{i|RW}}))$ as sampled from the joint posterior for parameters in our statistical model. A logarithmic change of zero ($\ln(1) = 0$; solid vertical line) represents no change over time suggesting that populations are at/near their equilibrium. The top row shows the predictions based on the community-level grand mean or the 'average' species. The second row shows the predicted change in percent cover for all species at this site illustrating how species vary within and among treatments. Treatment models did not receive any model support at this site so their predictions reflect the ambient predictions. Since this metric includes observed previous cover, differences seen in unsupported treatments reflect variations in species' cover rather than inferred parameters.

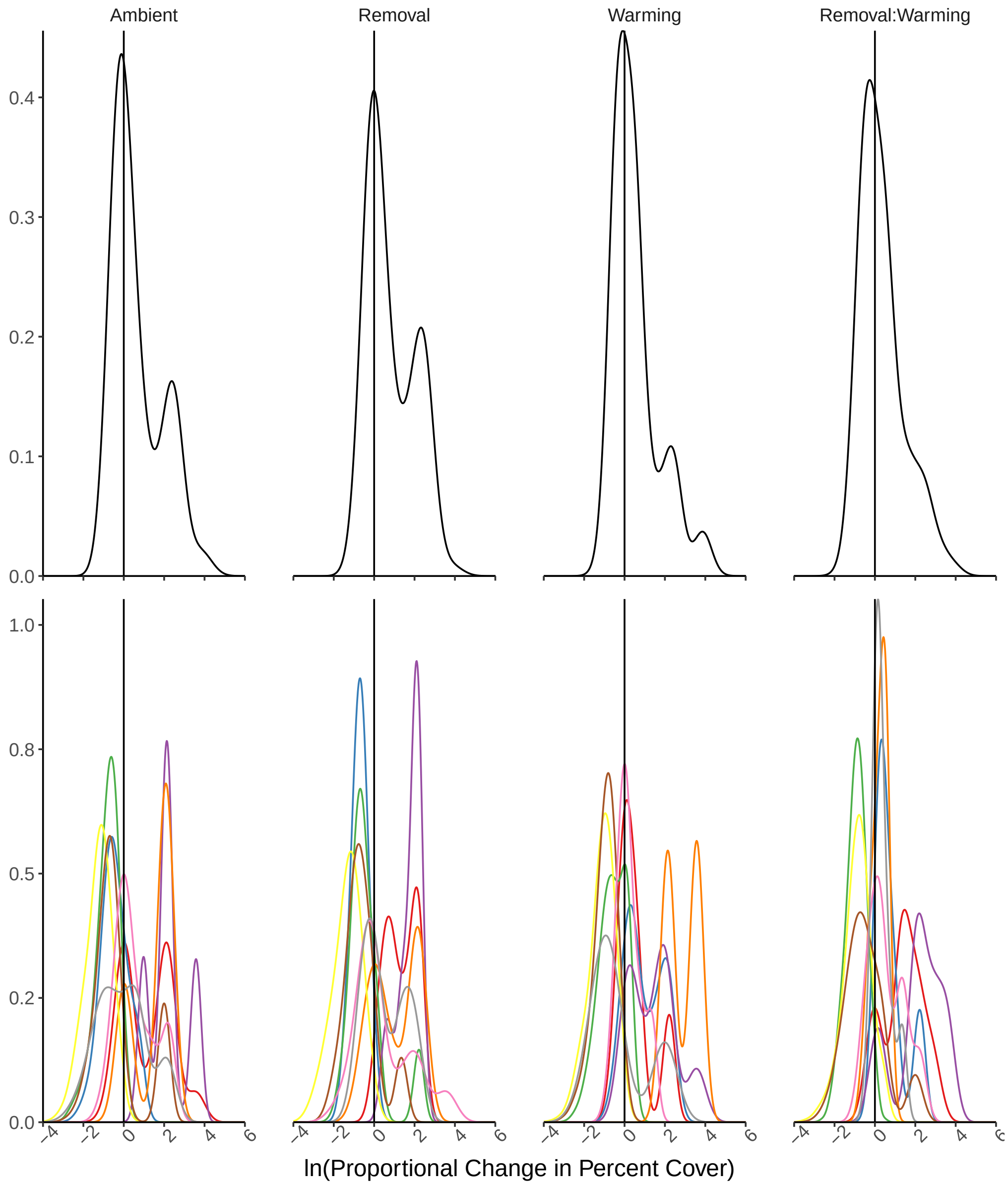


Fig. S12: Distribution of predicted change in percent cover within the high elevation New Zealand site. Logarithmic proportional change in cover is calculated as $(\ln(N_{i,t}/N_{i,t-1}) = \ln(Q_i/N_{i,t-1} + e^{G_{i|RW}}))$ as sampled from the joint posterior for parameters in our statistical model. A logarithmic change of zero ($\ln(1) = 0$; solid vertical line) represents no change over time suggesting that populations are at/near their equilibrium. The top row shows the predictions based on the community-level grand mean or the 'average' species, which in this case only shows minor effects of the removal treatment. The second row shows the predicted change in percent cover for all species at this site illustrating how species vary within and among treatments. This site showed support for multiple models and the posterior predictions were averaged across winning models: Removal × Warming, Warming, and Removal + Warming. Since this metric includes observed previous cover, differences seen in unsupported treatments reflect variations in species' cover rather than inferred parameters.

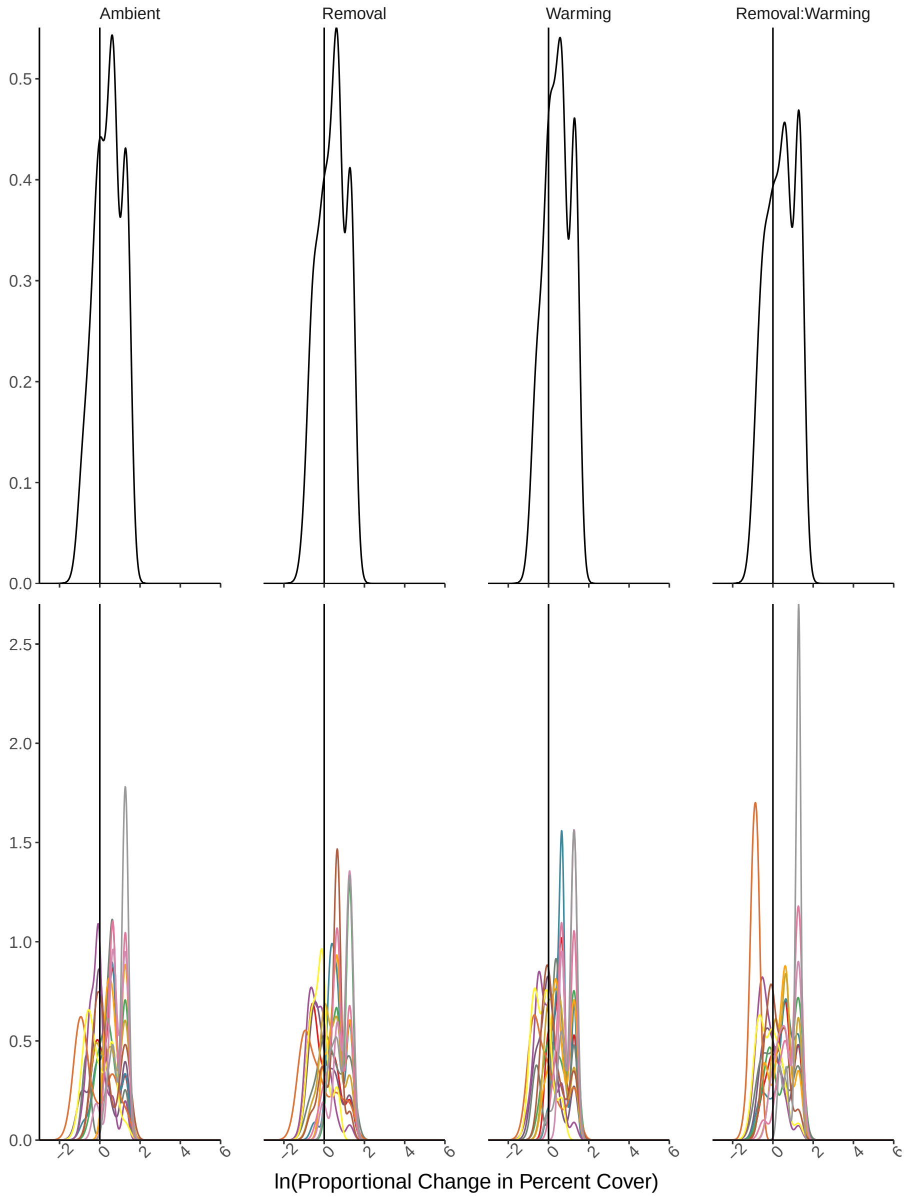


Fig. S13: Distribution of predicted change in percent cover within the low elevation United States site. Logarithmic proportional change in percent cover is calculated as a combination of density-independent and density-dependent factors ($\ln(N_{i,t}/N_{i,t-1}) = \ln(Q_i/N_{i,t-1} + e^{G_{i|RW}})$) is sampled from the posterior for each parameter in the model. A change in cover near zero represents no change over time suggesting that populations are at/near their equilibrium. The top row shows the predictions based on the community-level grand mean or the 'average' species. The second row shows the predicted change in percent cover for all species at this site illustrating how species vary within and among treatments. Treatment and combination models did not receive any model support at this site so their predictions reflect the ambient predictions. Since this metric includes observed previous cover, differences seen in unsupported treatments reflect variations in species' cover rather than inferred parameters.

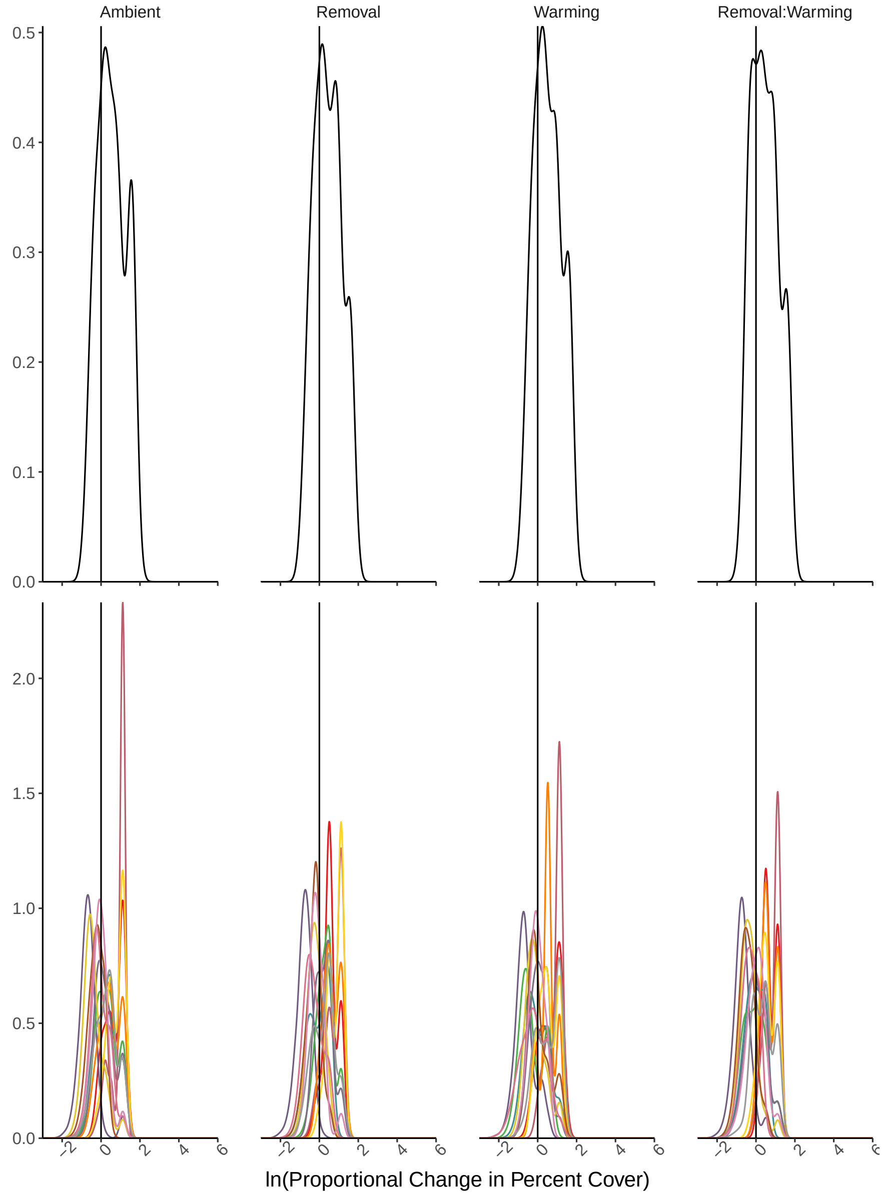


Fig. S14: Distribution of predicted change in percent cover within the high elevation United States site. Logarithmic proportional change in percent cover is calculated as a combination of density-independent and density-dependent factors ($\ln(N_{i,t}/N_{i,t-1}) = \ln(Q_i/N_{i,t-1} + e^{G_{i|RW}})$) is sampled from the posterior for each parameter in the model. A change in cover near zero represents no change over time suggesting that populations are at/near their equilibrium. The top row shows the predictions based on the community-level grand mean or the 'average' species, which in this case only shows minor effects of the warming treatment. The second row shows the predicted change in percent cover for all species at this site illustrating how species vary within and among treatments. This site showed support for multiple models and the posterior predictions were averaged across winning models: Warming, Ambient, and Removal + Warming. Since this metric includes observed previous cover, differences seen in unsupported treatments reflect variations in species' cover rather than inferred parameters.
